# Therapeutic targeting of MYC- and MYCN-driven medulloblastoma with a novel MYC degrader molecule

**DOI:** 10.64898/2026.07.09.737604

**Authors:** Sin Wi Ng, Satyanarayana Gadde, Nga-yee Chung, Qian Wang, Larissa Doughty, Tracy L Nero, Nisitha Jayatilleke, Janith Seneviratne, Daniel R Carter, Marion K Mateos, Maria Tsoli, David S. Ziegler, Raelene Endersby, Naresh Kumar, Louis Chesler, Tao Liu, Michael W Parker, Belamy B Cheung, Glenn M Marshall

## Abstract

**Background:** Medulloblastoma (MB) is the most common malignant brain tumour in children, and aggressive subgroups are frequently driven by the oncoproteins MYC or MYCN. Direct therapeutic targeting of MYC/MYCN has been challenging because of their intrinsically disordered protein structures. The aim of this study was to determine whether novel SE486-11 analogues (UNSW-SCs) can therapeutically target MYC/MYCN-driven MB.

**Methods:** The anticancer activity of UNSW-SCs was assessed in MB cell lines with differential MYC/MYCN expression. Target engagement was evaluated using surface plasmon resonance and drug affinity responsive target stability assays. Blood-brain barrier penetration, MYC/MYCN protein degradation, cell cycle effects, apoptosis, DNA damage, and synergy with histone deacetylase (HDAC) inhibitors were examined. Therapeutic efficacy was evaluated in murine models of MYC- and MYCN-driven human MB.

**Results:** UNSW-SCs showed potent anticancer activity, with preferential selectivity toward MB cells expressing high MYC/MYCN levels and IC50 values ranging from 0.22 to 1.18 μM. The lead molecule, UNSW-SC-22, directly bound MYC, crossed the blood-brain barrier, and achieved a brain-to-plasma ratio of 1.44 at peak concentrations. UNSW-SC-22 induced MYC/MYCN-dependent cytotoxicity associated with enhanced proteasomal degradation, cell cycle arrest, apoptosis, and DNA damage. Combined treatment with HDAC inhibitors further reduced MYC/MYCN protein levels, increased DNA damage, and enhanced apoptosis. *In vivo*, UNSW-SC-22, either alone or with entinostat, significantly suppressed intracranial tumour growth and prolonged survival.

**Conclusions:** UNSW-SC-22 is a brain-penetrant MYC/MYCN-targeting molecule with potent preclinical activity in MYC/MYCN-driven MB, supporting its development as a monotherapy or combination strategy with HDAC inhibition.

**Key Points:**

- UNSW-SC-22 directly targets MYC/MYCN in medulloblastoma.
- UNSW-SC-22 crosses the blood-brain barrier and prolongs survival.
- HDAC inhibition enhances UNSW-SC-22 activity in MYC-driven cells.

**Importance of the Study:** MYC- and MYCN-driven medulloblastomas remain among the most aggressive paediatric brain tumours, yet direct pharmacological targeting of MYC/MYCN has historically been difficult. This study identifies UNSW-SC-22 as a novel brain-penetrant small molecule that directly engages MYC/MYCN and promotes proteasomal degradation, leading to tumour cell death. Compared with prior MYC-directed approaches, UNSW-SC-22 combines direct target engagement, blood-brain barrier penetration, and *in vivo* efficacy in orthotopic MYC- and MYCN-driven medulloblastoma models. The study further demonstrates that combining UNSW-SC-22 with histone deacetylase inhibition enhances MYC/MYCN suppression, DNA damage, apoptosis, and survival benefit. These findings provide a translational framework for developing MYC/MYCN degradation as a therapeutic strategy for high-risk medulloblastoma and support future pharmacological optimisation, biomarker-guided patient selection, and combination therapy development.

## INTRODUCTION

Although outcomes for MB patients have improved over the past three decades, children with high-risk disease continue to respond poorly to conventional chemo-radiotherapy, with mortality rates of up to 70% ^1^. These high-risk cases are frequently associated with overexpression or amplification of the MYC oncogene ^2,3^.

MYC overexpression is a key driver of malignancy and is observed in 60-70% of all human cancers ^4,5^. While MYC has pleiotropic transcriptional effects, its binding at super-enhancers is thought to amplify survival signals in cancer cells. Although normal cells require transient MYC induction to initiate replication, preclinical and emerging clinical evidence suggests that short-term MYC inhibition has limited lasting effects on normal tissues ^6,7^. Together, these observations support MYC as a compelling therapeutic target in cancer. Although MYC is largely an intrinsically disordered protein, several MYC-targeted inhibitors have been developed ^8,9^. However, to date, only Omo-myc, a MYC dominant negative miniprotein, and the novel small molecule WBC100 (14-D-Valine-TPL), have advanced to early phase clinical trials ^10,11^. Moreover, the use of targeted agents in brain cancer has been severely limited by poor blood-brain barrier (BBB) penetration and tumour specificity.

Histone deacetylases (HDACs) are transcriptional repressors frequently dysregulated in cancer and can cooperate with MYC to regulate genes and pathways that drive oncogenic processes ^12–15^. Though there are several FDA-approved HDAC inhibitors used in cancer therapy, their clinical use is often limited by high toxicity and modest efficacy as monotherapies ^16^. Through an unbiased small molecule screen, we identified a novel pyrido-benzimidazole analogue, SE486-11, which enhanced the antitumour effects of the HDAC inhibitor SAHA in a panel of MYCN-amplified neuroblastoma (NB) cell lines ^17^. Unexpectedly, SE486-11 markedly reduced MYCN stability. We subsequently demonstrated that the SE486-11 analogue, (hereafter referred to as UNSW-SC-1), exhibited potent cytotoxic activity against NB cell lines in a MYCN-dependent manner, and synergy with HDAC inhibitors ^18^. We generated multiple UNSW-SC-1 analogues through structure-based design to improve efficacy. Given the high structural homology between MYCN and MYC, we investigated the anticancer effects of these analogues in MYC-driven MB. UNSW-SC-22 showed MYC-dependent cytotoxicity across MB cell lines and significantly reduced tumour burden in MYC- and MYCN-driven MB mouse models. Mechanistically, UNSW-SC-22 directly bound MYC and reduced its stability through increased Thr58 phosphorylation and proteasome-dependent degradation. These findings support UNSW-SC-22 as a novel MYC degrader with therapeutic potential as a monotherapy or combination strategy for high-risk MYC-driven MB.

## METHODS AND MATERIALS

### Cell Lines and Neurospheres Tissue Culture

Human MB cell lines D283MED, D341MED, CHLA-01-MED and CHLA-01R-MED, and human fibroblast lines MRC-5 and WI-38, were purchased from ATCC. D425MED was purchased from Merck. Normal human astrocytes were obtained from Lonza, and P000302 astrocytes were supplied by the ZERO team at CCI Australia. DAOY, UW228 and P493-6 Burkitt’s lymphoma cells were obtained from CCIA. GTML/Trp53^KI^/^KI^ MB neurospheres were kindly provided by Prof Louis Chesler, Institute of Cancer Research, London. DAOY, UW228 and D425MED cells were cultured in DMEM with 10% FBS. D283MED and D341MED were cultured in EMEM with 15% FBS. CHLA-01-MED and CHLA-01R-MED were maintained in DMEM/F12 supplemented with 20 ng/mL EGF, 20 ng/mL bFGF and 2% B-27. MRC-5 and WI-38 cells were cultured in MEMα with 10% FBS, while NHA and P000302 astrocytes were cultured in Astrocyte Growth Medium BulletKit. GTML/Trp53^KI^/^KI^ neurospheres were maintained in Neurobasal-A medium supplemented with 20 ng/mL EGF, 20 ng/mL bFGF and 2% B-27 without vitamin A. P493-6 cells were cultured in RPMI-1640 with 10% FBS. GTML/Trp53^KI^/^KI^ neurospheres and P493-6 cells express MYCN and MYC, respectively, under tetracycline-regulated promoters, with doxycycline inducing MYCN/MYC downregulation. All cells were cultured at 37°C with 5% CO₂, maintained in exponential growth phase, passaged as required and routinely tested for mycoplasma contamination.

### Cell Viability Assays and Drug Synergy Calculations

Cells were seeded in 96-well plates at densities optimised to reach ∼90% confluence at assay endpoint. Adherent cells were allowed to attach for 24 h before treatment. UNSW-SC compounds were synthesised by Dr Satyanarayana Gadde at School of Chemistry, UNSW Sydney; all other drugs were commercially sourced. For single-agent assays, cells were treated across appropriate dose ranges for 72 h at 37°C with 5% CO₂. Cell viability was assessed using resazurin, with fluorescence measured at 560/590 nm on Victor3 or Varioskan LUX microplate readers before and after 5-6 h incubation. Background-subtracted values were normalised to vehicle controls. IC₅₀ values were calculated in GraphPad Prism 9 using nonlinear regression. Data are presented as mean ± SEM from at least three independent biological replicates. For combination treatments, a 6×5 matrix drug screen was performed, whereby the concentrations used for each combination were values flanking the IC_50_ values of each drug, such as ¼ x IC_50_, ½ x IC_50_, 1 x IC_50_, 2 x IC_50_ and 4 x IC_50_. Drug synergy was calculated with excess over the highest single agent (HSA) model ^19^ and the BLISS model of drug independence (BLISS) ^20^ to generate synergism indices (SI). An SI > 0 indicates synergism, SI = 0 indicates additivity and SI < 0 indicates antagonism, and is obtained from at least three independent biological replicates.

### Generation of MYC-Inducible Cell Lines

Stable MB cell lines were generated from high MYC MB cell lines D283MED and D425MED using doxycycline-inducible GFP-IRES-shRNA FH1tUTG lentiviral constructs with short hairpin RNA (shRNA) sequences targeting a scrambled control (shCTRL: GCACTACCAGAGCTAACTCAGATAGTACT) or MYC (shMYC2: GATGAGGAAGAAATCGATG). The shRNA sequences were cloned into the lentiviral construct, expanded in *E.coli* competent cells and transfected using Lipofectamine 2000 into HEK293T cells for viral production. The collected viral media was then concentrated and transduced with polybrene into D283MED and D425MED MB cell lines and sorted for the top 10% of cells expressing GFP. Validation of MYC knockdown was conducted via western blot.

### Western Blotting

Whole-cell and mouse brain tissue lysates were prepared in RIPA buffer containing protease inhibitors, with phosphatase inhibitors added when phosphorylated proteins were analysed. Brain tissues were homogenised using a TissueRuptor. Lysates were incubated on ice for 30 min, cleared by centrifugation, and protein concentrations were determined using the Pierce BCA assay. Proteins (30-40 µg) were denatured in loading buffer containing DTT, separated on Criterion gels and transferred to nitrocellulose membranes. Membranes were stained with Ponceau S, blocked in 10% skim milk/TBS-T for 1 h, and incubated overnight at 4°C with primary antibodies against c-MYC, MYCN, MELK, USP1, PARP, caspase/cleaved caspase-3, p21, H2AX, phospho-H2AX, p53, vinculin, GAPDH or β-actin. After washing, membranes were incubated with goat anti-rabbit or goat anti-mouse secondary antibodies for 2 h at room temperature. Proteins were detected using Clarity Western ECL substrate and imaged on a ChemiDoc Touch system. Vinculin, GAPDH or β-actin were used as loading controls. Band intensities were quantified using Quantity One software and normalised to loading and experimental controls. Data were derived from at least three independent biological replicates. Antibodies used are detailed in **Table 1**.

### Quantitative Real-Time Polymerase Chain Reaction (qRT-PCR)

Total RNA was extracted using the PureLink RNA Mini Kit (ThermoFisher) and quantified with a NanoDrop 1000 (ThermoFisher). cDNA was synthesised from 1 μg RNA using the Tetro cDNA Synthesis Kit (Bioline). qRT-PCR was performed using Power SYBR Green master mix with forward and reverse primers in MicroAmp Optical 96- or 384-well plates on a QuantStudio 3 or 5 system (ThermoFisher). Cycling conditions were 90°C for 3 minutes, followed by 40 cycles of 95°C for 15 seconds and 60°C for 1 minute. Gene expression was normalised to B2M and respective controls using the ΔΔCt method. Primer sequences are listed in **Table 2**.

### Patient Tumour Cohort and Cell Line Analyses

Microarray data (GSE85217) of 549 clinically annotated MB samples ^21^ were obtained from GEO. Gene expression levels of MYC were extracted and dichotomised at the upper quartile as they had a left-skewed distribution. For genes identified from RNA-seq analysis of D283MED cells, expression levels were dichotomised by median expression. Multivariate Cox proportional hazards models were fitted for each gene using the survival R package to assess associations with overall survival. Kaplan–Meier survival curves were generated using survminer. Bulk RNA-seq data for medulloblastoma and normal human astrocyte cell lines were obtained from GEO (GSE185024, GSE171117, GSE52983, GSE102505) and ArrayExpress (MTAB-9833). Raw MYC counts were extracted and normalised across cell lines using DESeq2 v1.34.0. Gene expression differences were visualised using ggplot2.

### Drug Affinity Responsive Targets Stability (DARTS) Assay

DARTS assay was conducted in D283MED cells according to the previously published protocol ^22^. Cells were lysed with M-PER buffer containing phosphatase and protease inhibitors. Lysates were incubated with DMSO or UNSW-SC-22 for 15 minutes at room temperature, followed by Pronase digestion for 30 minutes. Reactions were quenched with protease inhibitor and analysed by western blot. Densitometry was normalised to Vinculin and then to the DMSO control.

### Surface Plasmon Resonance (SPR)

SPR experiments were performed on a Biacore T200 biosensor (Cytiva) using a CMD500M chip (Xantec). MYC protein was immobilised in HBS-EP buffer following chip activation with NHS/EDC and blocked with ethanolamine, achieving immobilisation levels of ∼7,000 RU. Binding assays were conducted in HBS-EP containing 5% DMSO. UNSW-SC-22 was serially diluted two-fold from 5 µM, injected for 70 seconds, and dissociated for 90 seconds at 30 µL/min. Sensorgrams were solvent-corrected, double-referenced and analysed using Biacore T200 Evaluation Software v3.2.1. Steady-state affinities were determined using a 1:1 binding model. Data were obtained from three independent biological replicates.

### Medulloblastoma Orthotopic Xenografts

Balb/c nude mice aged 6-7 weeks were intracranially injected with 50,000 D425MED-luciferase medulloblastoma cells in Matrigel into the cerebellum using a stereotactic device (Kopf Instruments; coordinates: 2 mm lateral, 2 mm posterior and 2 mm deep relative to Lambda). Mice were anaesthetised with 2.5% isoflurane in oxygen during the procedure and monitored daily for 7 days for weight loss, dehydration and neurological symptoms. Mice were randomised 10 days after injection and treated intraperitoneally with DMSO vehicle or UNSW-SC-22 at 90 mg/kg/day on a 5-days-on, 2-days-off schedule for 21 days. Tumour growth was monitored weekly by bioluminescence imaging, and mice were monitored daily for weight loss and neurological symptoms. Mice showing >20% weight loss or severe neurological symptoms were humanely euthanised with CO₂. At endpoint, brains were collected and fixed in 10% neutral buffered formalin for histological analysis. Kaplan-Meier survival curves were generated in GraphPad Prism 9 and compared using two-sided log-rank tests.

### Transgenic GTML/p53ER mice

A GTML/p53ER (GTML; *Glt1-tTA/TRE-MYCN-LUC*, p53; *Trp35^KI/KI^)* mice colony was generated in-house through crossing of 2 different transgenic mouse models; (i) a doxycycline-regulable MYCN-driven mouse model (GTML, Glt1-tTA/TRE-MYCN-LUC), and (ii) a tamoxifen-regulable p53 deficient mouse model (p53ER; Trp53KI/KI) ^23,24^. Homozygous GTML/p53ER mice developed tumours at 3-4 weeks of age with 100% incidence. Following tumour detection by bioluminescence imaging, mice were randomised and treated intraperitoneally with DMSO vehicle, UNSW-SC-22 at 90 mg/kg/day, or entinostat at 10 mg/kg/day on a 5-days-on, 2-days-off schedule for 21 days. Tumour growth was monitored weekly by bioluminescence imaging, and mice were assessed daily for weight loss and neurological symptoms. At endpoint, brains were collected and fixed in 10% neutral buffered formalin for histological analysis.

### Statistical Analyses

All experiments performed included a minimum of two independent biological replicates. All statistical analyses were performed using GraphPad Prism 9, and p-values were calculated using one-way ANOVA, two-way ANOVA or unpaired t-tests. Error bars represent the standard error of the mean (SEM) between replicates. R (v4.1.2) and R Studio (v2021.09.0) were also used.

## RESULTS

### UNSW-SC Compounds Exhibit Potent and Selective Anticancer Activity Against Medulloblastoma Cells

We previously identified the small molecule SE486-11 through a high-throughput screen for compounds that enhanced the cytotoxic effect of SAHA on the cell viability of NB cells ^17^. Unexpectedly, SE486-11 also markedly reduced MYCN protein stability. To improve potency, we synthesised a series of SE486-11 analogues, collectively termed UNSW-SC compounds (**Fig. 1A**), and evaluated their efficacy as monotherapies in MB models. A panel of seven molecularly heterogeneous human MB cell lines was treated with four UNSW-SC compounds: UNSW-SC-1, -22, -36 and -61. These were selected based on their previously demonstrated potency in MYCN-driven neuroblastoma cells. UNSW-SC-1, -22 and -61 were tested at 0-20 μM (**Fig. 1B**), while the more potent UNSW-SC-36 was tested at 0-0.3125 μM (**Fig. 1C**). Cell viability was assessed after 72 hours relative to vehicle control. All four compounds showed dose-dependent cytotoxicity across MB cell lines, with IC₅₀ values ranging from 0.017 to 6.88 μM. To assess selectivity, three normal cell lines, including one fibroblast and two human astrocyte lines, were treated with UNSW-SC-1, -22 and -61 at similar dose ranges **(Fig. S1)**. These compounds showed substantially reduced cytotoxicity in normal cells, with IC₅₀ values ranging from 2.18 to >10 μM (**Fig. 1D**). By contrast, UNSW-SC-36 showed limited selectivity, with IC₅₀ values of 0.05 to >0.3 μM in normal cells compared with 0.017–0.04 μM in MB cells. Together, these findings indicate that UNSW-SC-1, -22 and -61 selectively target MB cells, supporting their potential as therapeutics for high-risk MB.

**Figure 1.**
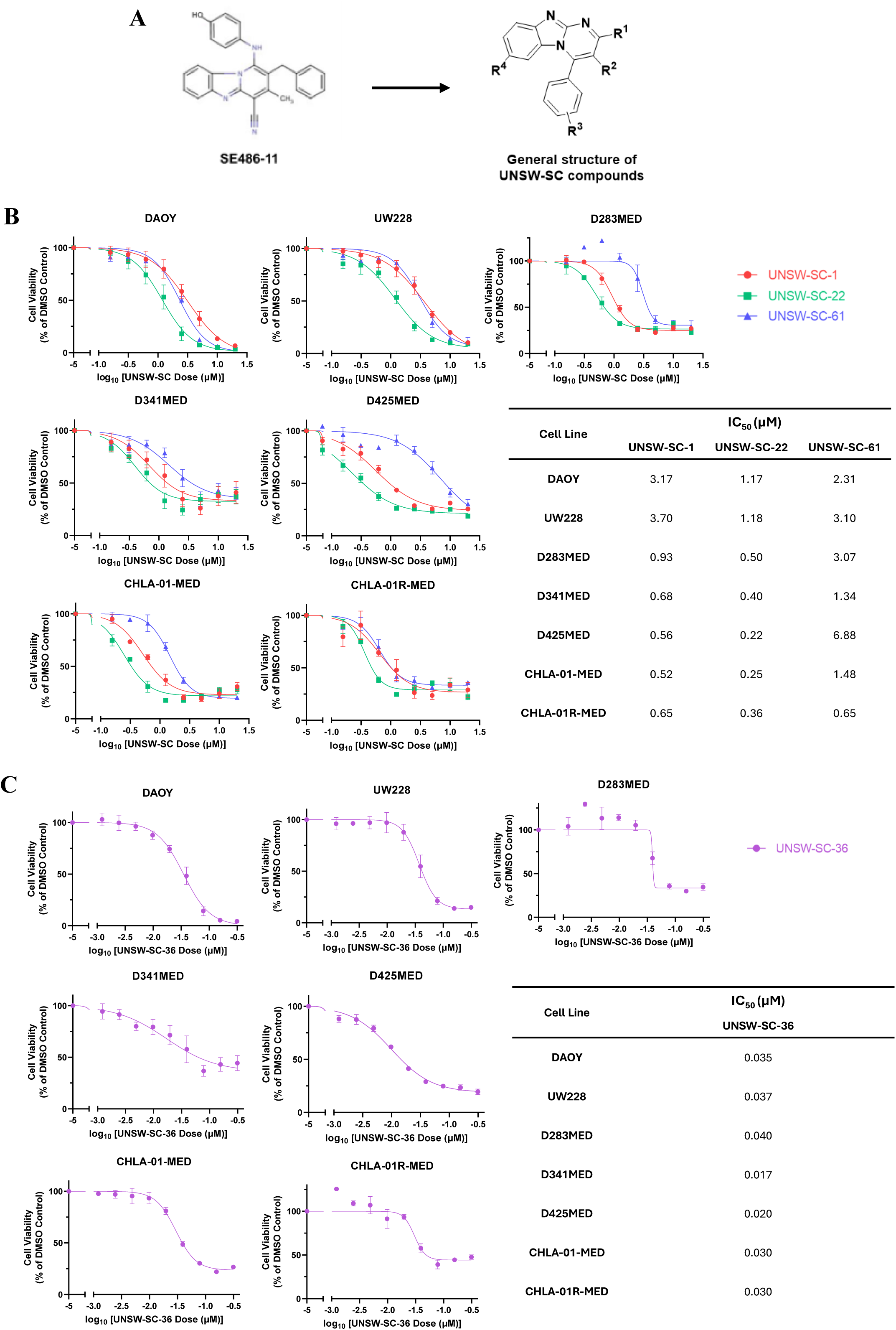
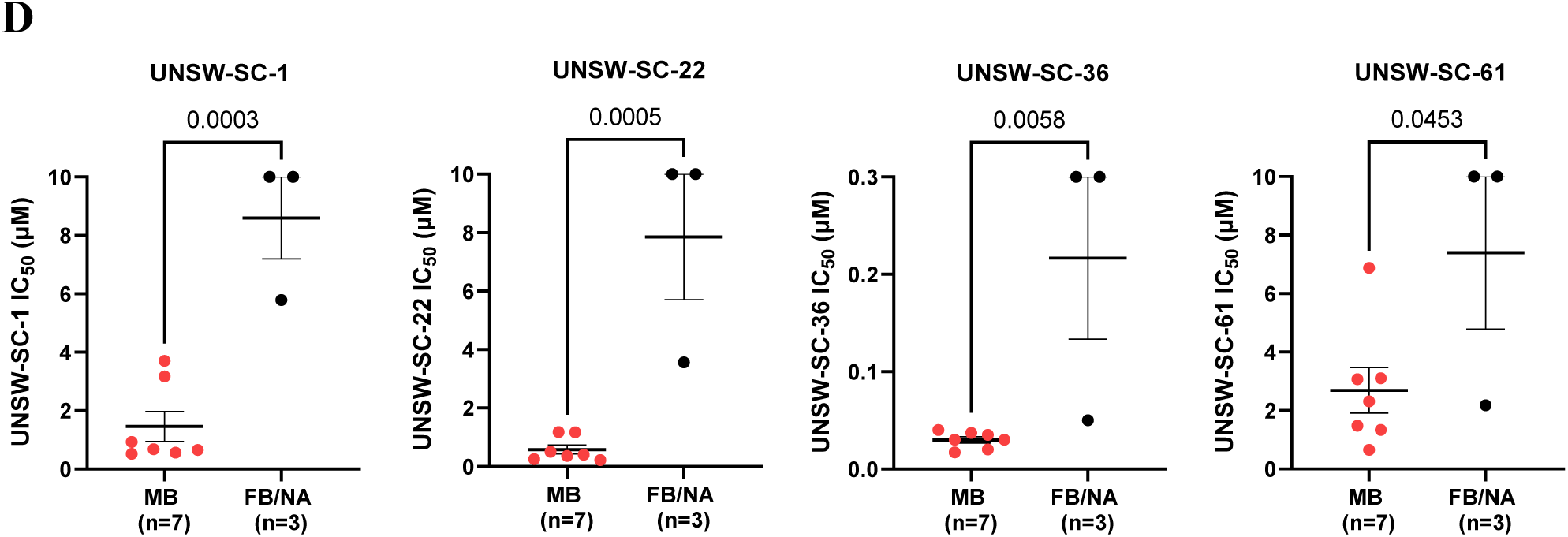
UNSW-SC compounds exhibit potent and specific anti-cancer activity against medulloblastoma cells. **(A)** The chemical structure of SE486-11, and the general structure of UNSW-SC compounds. **(B)** Cell viability curves and table of 7 MB cell lines upon treatment with UNSW-SC-1, UNSW-SC-22, and UNSW-SC-61 (0 – 20 μM) for 72 hours. Values represent cell viability relative to a DMSO vehicle control. **(C)** Cell viability curves of 7 MB cell lines upon treatment with UNSW-SC-36 (0 – 0.3125 μM) for 72 hours. **(D)** Comparison of IC_50_ of 4 UNSW-SC compounds in 7 medulloblastoma (MB) cell lines, 1 fibroblast cell line and 2 normal human astrocytes (FB/NA). Each point represents average cell viability or IC_50_ values across atleast three independent experiments. Error bars represent the standard error of the mean and statistical comparisons were performed using a t-test.

### MYC Expression is a Key Determinant Of Sensitivity to UNSW-SC Compounds

We hypothesised that MB cell sensitivity to UNSW-SC compounds is determined by MYC expression. To test this, we analysed compound cytotoxicity (IC₅₀) in relation to MYC mRNA expression using publicly available RNA-seq datasets (**Fig. S2A**), and MYC protein levels by immunoblotting in corresponding MB and normal cell lines (**Fig. 2A**). Based on MYC expression, MB cell lines were stratified into MYC-high (D283MED, D341MED, D425MED, CHLA-01-MED and CHLA-01R-MED) and MYC-low (DAOY and UW228) groups. MYC-high cell lines were, on average, 5.14-fold and 3.40-fold more sensitive to UNSW-SC-1 and UNSW-SC-22, respectively, than MYC-low lines (**Fig. 2B**). In contrast, no significant difference was observed for UNSW-SC-36 or UNSW-SC-61 (Fig. 2B). IC₅₀ values showed a strong negative correlation with MYC protein levels for UNSW-SC-1 (r = −0.8775, p = 0.0216) and UNSW-SC-22 (r = −0.8188, p = 0.0463) (**Fig. 2C**), but not for UNSW-SC-36 or UNSW-SC-61 (**Fig. S2B**). Similar correlations were observed between MYC RNA expression and sensitivity to UNSW-SC-1 and -22 (**Fig. S2C**). Given its favourable therapeutic index, potent activity across MB cells (IC₅₀ 0.22-1.18 μM), strong association with MYC expression, and selective cytotoxicity in MYC-high MB cells, UNSW-SC-22 was selected for further evaluation of efficacy, specificity and mechanism of action.

**Figure 2.**
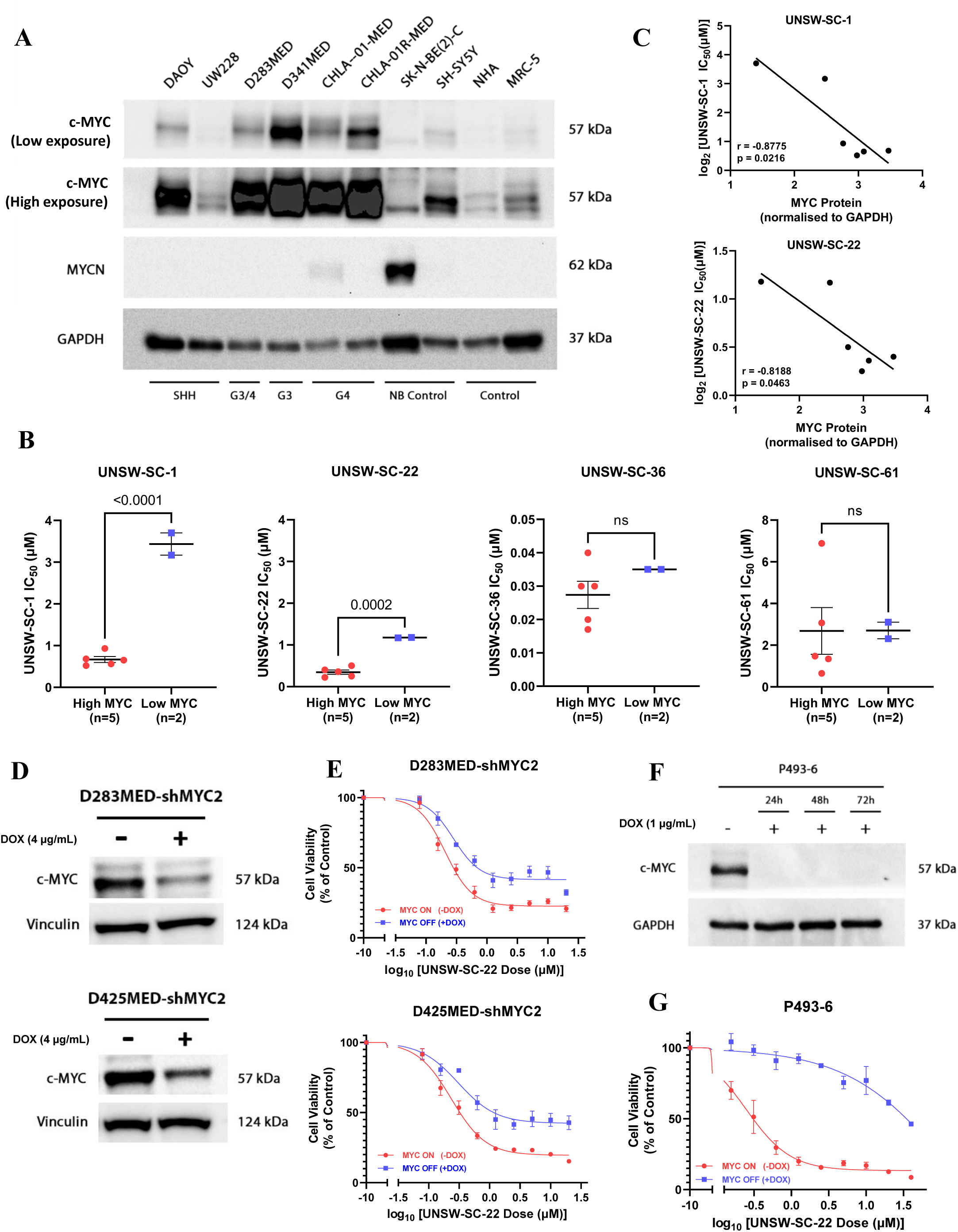
MYC expression is a key determinant of UNSW-SC-1 and UNSW-SC-22 drug sensitivity. **(A)** Immunoblot of MYC and MYCN proteins in MB, neuroblastoma, NA and FB cell lines. **(B)** IC_50_ comparison for UNSW-SC-1, UNSW-SC-22, UNSW-SC-36 and UNSW-SC-61 in 7 MB cell lines based on MYC status. **(C)** Pearson correlation between MYC protein levels (from Figure 2A) and IC_50_ of UNSW-SC-1 and UNSW-SC-22. **(D)** Immunoblot of MYC proteins in D283MED-shMYC2 and D425MED-shMYC2 MB cells in the presence or absence of 4μg/mL of doxycycline for 72 hours. **(E)** Cell viability curves of D283MED-shMYC2 and D425MED-shMYC2 MB cell lines upon treatment with UNSW-SC-22 (0 – 20 μM) for 72 hours in the presence or absence of 4μg/mL of doxycycline. **(F)** Immunoblot of MYC protein in P493-6 Burkitt’s lymphoma cells in the presence or absence of 1μg/μL of doxycycline for 72 hours. **(G)** Cell viability curves of P493-6 Burkitt’s lymphoma cells upon treatment with UNSW-SC-22 (0 – 20 μM) for 72 hours in the presence or absence of 1μg/μL of doxycycline. Values represent cell viability relative to a DMSO vehicle control. Each point represents average cell viability or IC_50_ values across at least three independent experiments. Error bars represent the standard error of the mean and statistical comparisons were performed using one-way ANOVA, two-way ANOVA, or t-test.

To further validate MYC as the molecular target of UNSW-SC-22, we generated doxycycline-inducible MYC shRNA knockdown models in MYC-overexpressing MB cell lines D283MED and D425MED. Cells were treated with DMSO or doxycycline for 72 hours to induce MYC knockdown (**Fig. 2D**), followed by UNSW-SC-22 treatment at 0-20 μM for an additional 72 hours (**Fig. 2E**). MYC knockdown significantly reduced UNSW-SC-22 cytotoxicity, with sensitivity decreased by 2.96-fold in D283MED-shMYC2 cells (p = 0.0294) and 3.13-fold in D425MED-shMYC2 cells (p = 0.0429) (**Fig. 2E, Fig. S2D-F**). As MYC protein levels were only partially reduced in the shMYC2 models (∼50%), we also used the doxycycline-inducible MYC-expressing Burkitt’s lymphoma cell line P493-6, in which doxycycline fully represses MYC protein expression (**Fig. 2F**) ^25^. Treatment of P493-6 cells with the same UNSW-SC-22 dose range for 72 hours resulted in a >50-fold reduction in cytotoxic sensitivity following MYC repression (**Fig. 2G, Fig. S2G**). These findings provide strong evidence that MYC is a critical functional target of UNSW-SC-22.

### UNSW-SC-22 Reduces MYC Protein Levels and Downstream Transcriptional Targets MELK and USP1

Given the role of MYC as an oncogenic driver in MB, we examined MYC protein levels following UNSW-SC-22 treatment in two MB cell lines. UNSW-SC-22 significantly reduced MYC protein in a dose- and time-dependent manner (**Fig. 3A, Fig. S3A**). In contrast, MYC mRNA levels remained largely unchanged, with only a slight increase in CHLA-01R-MED cells at 24 hours, potentially reflecting an acute adaptive response, as MYC protein was substantially reduced by 72 hours (**Fig. S3B**). These findings suggest that UNSW-SC-22 reduces MYC primarily at the post-transcriptional level. To investigate the molecular mechanisms underlying UNSW-SC-22 cytotoxicity, we performed RNA-seq in MYC-overexpressing D283MED Group 3/4 MB cells treated with DMSO or 1.5 μM UNSW-SC-22 for 24 hours. This identified 512 differentially expressed genes, including 344 downregulated and 168 upregulated genes, using an absolute log₂ fold change >0.5 and adjusted p-value <0.05 (**Fig. 3B, Data file S1**). These genes were then analysed by GSEA using the HALLMARK gene sets from the Molecular Signatures Database (**Data file S1**) ^26^. This analysis showed that MYC targets, E2F targets and G2/M checkpoint gene sets were markedly downregulated following UNSW-SC-22 treatment (**Fig. 3C, Fig. S3C**). Conversely, p53 signalling was significantly enriched among upregulated transcripts (**Fig. 3C, Fig. S3C**). Consistent with this, UNSW-SC-22 treatment robustly increased p53 protein levels in two MYC-driven MB cell lines (**Fig. 3D**).

**Figure 3.**
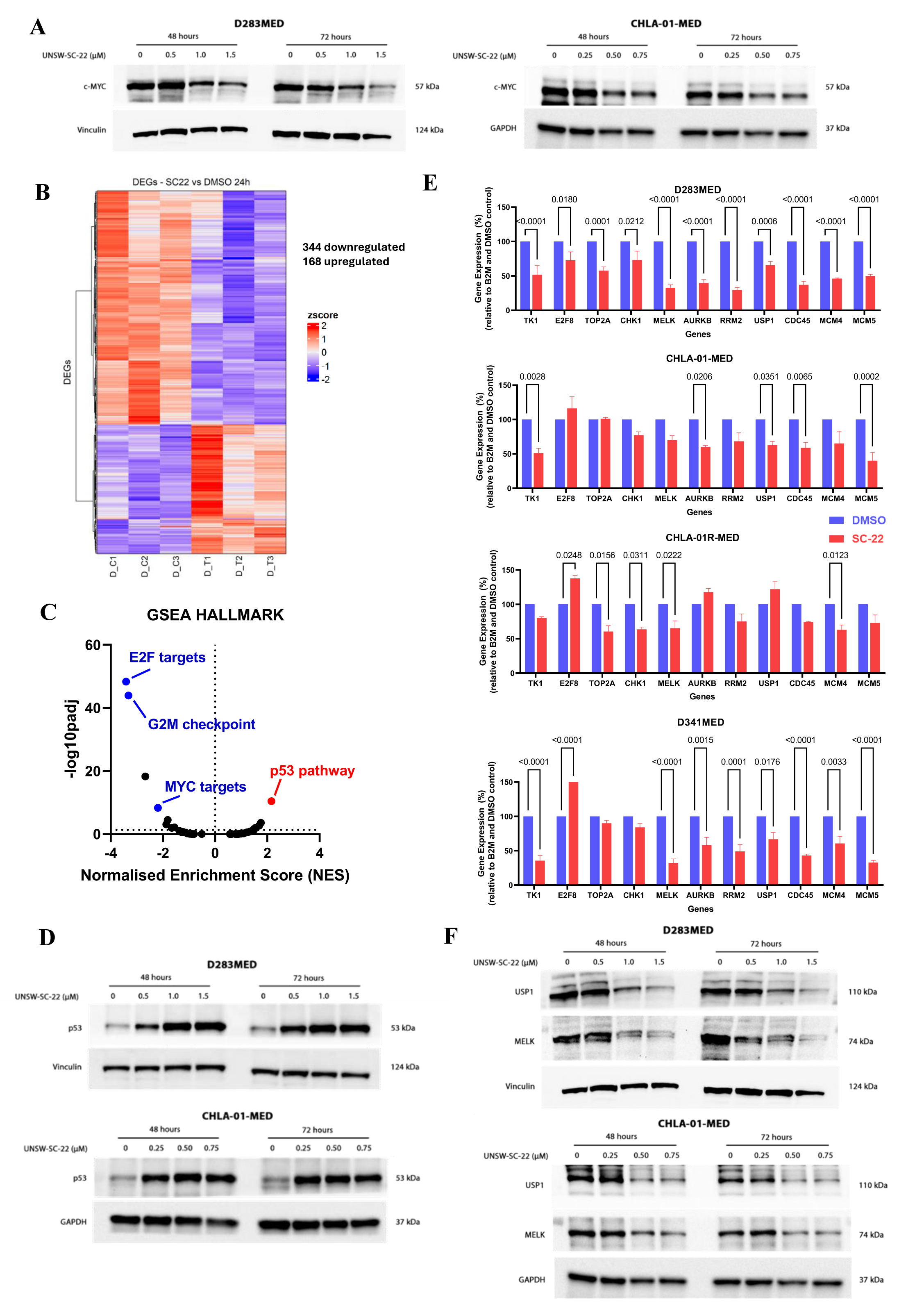
UNSW-SC-22 reduces MYC protein and downstream transcriptional factors MELK and USP1. **(A)** Immunoblot of MYC protein in the indicated treatment groups for D283MED and CHLA-01-MED MB cells. **(B)** Heatmap of 512 differentially expressed genes (DEGs) (344 downregulated genes, 168 upregulated genes) in group 3/4 D283MED cells treated with 1.5 μM of UNSW-SC-22 versus DMSO control for 24 hours. Each row represents z-score normalized gene expression across all samples. D_C1-3 indicates D283MED controls (DMSO) and D_T1-3 indicates D283MED treated with UNSW-SC-22. **(C)** Volcano plot of GSEA analysis of 512 differentially expressed genes with HALLMARK dataset from molecular signatures database. **(D)** Immunoblot of WT p53 protein in the indicated treatment groups for D283MED and CHLA-01-MED cell. **(E)** Real-time PCR analysis of mRNA expression for 11 selected genes following treatment of DMSO or 1.5 μM UNSW-SC-22 for 24 hours in high MYC expression MB cell lines D283MED, CHLA-01-MED, CHLA-01R-MED and D341MED cells. **(F)** Immunoblot of USP1 and MELK proteins in the indicated treatment groups for D283MED and CHLA-01-MED cells. Error bars represent the standard error of the mean and statistical comparisons were performed using two-way ANOVA. The loading controls in Figure 3A were reused in Figure 3D and 3F as samples were run on the same gel and are shown in separate panels for clarity.

To identify downstream targets mediating UNSW-SC-22 cytotoxicity, we selected 11 significantly downregulated genes for further validation by qRT-PCR in a panel of MB cell lines: E2F8, TOP2A, RRM2, AURKB, TK1, MELK, CDC45, MCM4, CHEK1, MCM5 and USP1 (p < 0.05, fold change > 1.5). Genes were selected based on their high expression in primary MB tumours and association with poor patient prognosis in the Cavalli dataset ^21^; (ii) a score of < 0 from the Cancer Dependency Map (DepMap) ^27^ indicating a level of gene dependency across MB cell lines and (iii) a known regulatory relationship between the gene and MYC **(Fig. S3, D and E).** We found a general decrease in expression of most genes across all 4 MB cell lines after treatment with 1.5 μM of UNSW-SC-22 for 24 hours, with the exception of E2F8 and TOP2A **(Fig. 3E)**. We then further looked at MELK and USP1, since both genes have been reported to be transcriptionally regulated by MYC at their promoter regions ^28,29^ **(Fig. S3F)**. Following UNSW-SC-22 treatment, MELK and USP1 protein levels decreased in a dose- and time-dependent manner (**Fig. 3F, Fig. S3G and H**). Collectively, these data suggest that UNSW-SC-22 targets MYC, reducing MYC protein levels and downstream MYC-regulated gene expression. We also assessed all 344 downregulated genes as a single gene signature, termed SC22DR. High SC22DR expression was associated with poorer overall survival in the Cavalli MB patient dataset (p = 0.004) (**Fig. S3I**), suggesting that UNSW-SC-22 suppresses a clinically relevant gene program linked to poor prognosis in MB.

### UNSW-SC-22 Directly Binds MYC and Promotes Its Proteasomal Degradation

To define the molecular basis of MYC inhibition, we assessed whether UNSW-SC-22 directly binds MYC. Surface plasmon resonance analysis showed dose-dependent binding between UNSW-SC-22 and MYC, with a micromolar affinity (K_D_ = 12.5 ± 3.0 µM) (**Fig. 4A–C**). In contrast, the inactive analogue UNSW-SC-24 showed no detectable MYC binding (**Fig. S4A–C**). To further validate this interaction, we performed a DARTS assay in D283MED cells. UNSW-SC-22 partially protected MYC from Pronase-mediated degradation (**Fig. 4D**), supporting direct target engagement between UNSW-SC-22 and MYC in MB cells.

**Figure 4.**
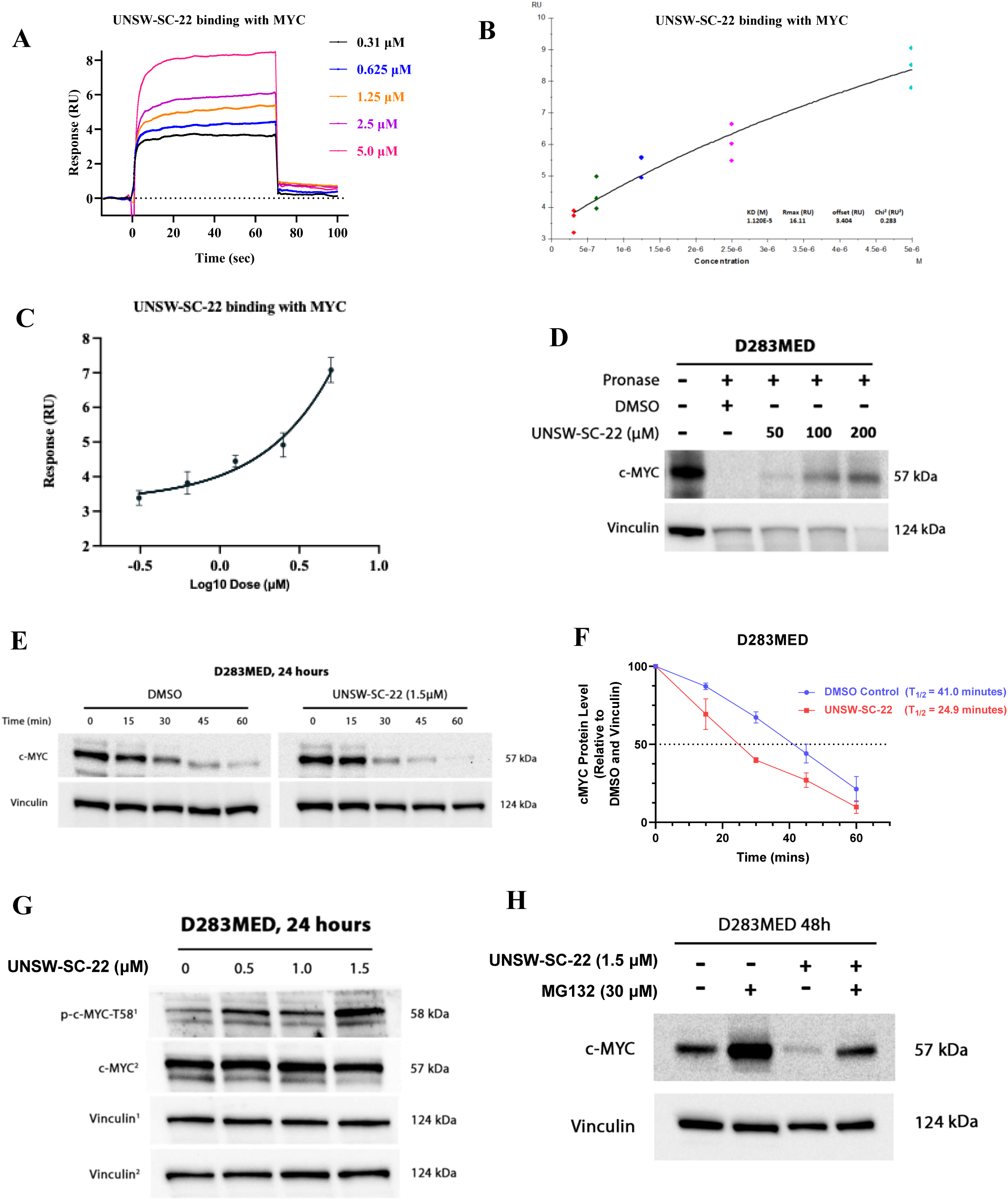
UNSW-SC-22 directly targets c-MYC and promotes its phosphorylation and ubiquitin-mediated proteasomal degradation. **(A-B)** Representative SPR curves illustrating the concentration-response binding of UNSW-SC-22 to MYC. **(C)** Normalized binding affinity curve showing a direct dose-response binding affinity between MYC and UNSW-SC-22. **(D)** Immunoblot of MYC gene in D283MED cell lysates after incubation with UNSW-SC-22 at the indicated doses, followed by digestion with pronase. **(E)** Immunoblot of MYC protein in the indicated treatment groups for D283MED cells following a 60-minute cycloheximide chase. **(F)** Densitometry analysis of Figure 4D, where MYC levels were first normalized to vinculin loading control and then to the 0-minute time point, error bars represent the standard error of the mean from three independent experiments. The half-life (t_1_/2) of MYC in each condition is reported. **(G)** Immunoblot of MYC-Thr58 and MYC protein at the indicated treatment groups for D283MED cells. **(H)** Immunoblot of MYC protein of D283MED cells treated with DMSO or 1.5 μM of UNSW-SC-22 for 48 hours, followed by incubation with or without 30μM of MG132 for 6 hours.

We next assessed whether UNSW-SC-22 reduced MYC protein stability using cycloheximide chase assays. D283MED cells were treated with DMSO or 1.5 μM UNSW-SC-22 for 24 hours, followed by 100 µg/mL cycloheximide for 0, 15, 30, 45 or 60 minutes. UNSW-SC-22 treatment reduced MYC protein half-life by 40% compared with DMSO control (**Fig. 4E and F**). UNSW-SC-22 also increased MYC phosphorylation at Thr58, a modification associated with reduced MYC stability and enhanced proteasomal degradation ^30,31^ **(Fig. 4G, Fig. S4D)**. To determine whether proteasomal degradation contributed to increased MYC turnover, D283MED cells were treated with DMSO or UNSW-SC-22 for 48 hours, followed by the proteasome inhibitor MG-132 at 30 μM for 6 hours. MG-132 markedly increased MYC protein levels in DMSO-treated cells, whereas this increase was blocked following UNSW-SC-22 treatment (**Fig. 4H, Fig. S4E**). Overall, these data suggest that UNSW-SC-22 directly interacts with MYC, promoting its destabilisation and proteasomal degradation.

### UNSW-SC-22 Induces Cell Cycle Arrest, Apoptosis and DNA Damage in MB Cells

We observed that a small population of MB cells remained viable despite logarithmic dose escalation of UNSW-SC compounds, suggesting a cytostatic effect. To investigate this, we assessed cell cycle dynamics following UNSW-SC-22 treatment using propidium iodide staining in D283MED cells, a MYC-overexpressing line representative of group 3 MB. The proportion of cells in G2/M phase increased almost two-fold at the highest UNSW-SC-22 concentration (p = 0.0007; **Fig. 5A and B, Fig. S5A to C**). This G2/M accumulation was dose-and time-dependent, suggesting that G2/M arrest contributes to the mechanism of action of UNSW-SC-22. Treatment also reduced the proportion of cells in S phase and increased G1-phase cells by 13–20% after 24 hours (p ≤ 0.0001), but not after 48 hours (**Fig. 5A and B, Fig. S5A to C**). A significant dose-dependent increase in the sub-G1 population was also observed (**Fig. 5C**). The marked increase in sub-G1 cells after 48 hours suggested DNA fragmentation and apoptosis. Consistent with this, Annexin V staining showed a significant dose-dependent increase in early and/or late apoptotic cells (**Fig. 5D, Fig. S5D**). UNSW-SC-22 treatment also induced p21 expression and promoted dose-dependent cleavage of PARP-1 and caspase-3 (**Fig. 5E, Fig. S5E to G**). Given that sub-G1 accumulation can indicate DNA damage, we next assessed phosphorylated H2AX. UNSW-SC-22 treatment induced a dose-dependent increase in DNA damage, shown by increased phosphorylated H2AX expression compared with DMSO control (**Fig. 5F, Fig. S5H**). Together, these data demonstrate that UNSW-SC-22 exerts anticancer effects by inducing cell cycle arrest, apoptosis and DNA damage in MB cells.

**Figure 5.**
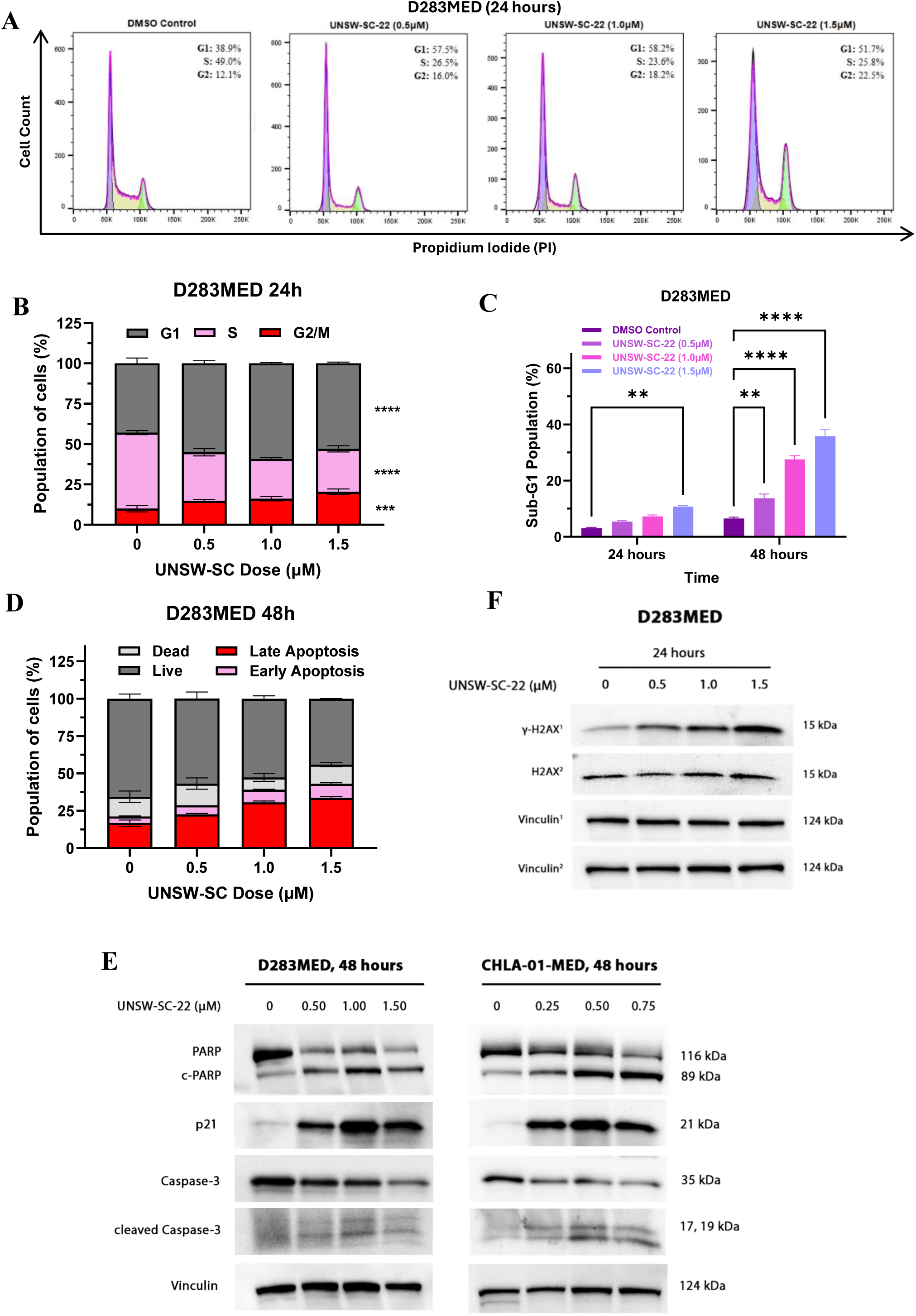
UNSW-SC-22 induces cell cycle arrest, apoptosis and DNA damage in medulloblastoma cell lines. **(A)** Flow cytometry analysis of cell cycle distribution in D283MED cells after treatment with UNSW-SC-22 at the indicated doses for 24 hours, followed by propidium iodide staining. **(B)** Cell cycle assay represented in stacked column graphs indicating the proportion of cells in each cell cycle phase 24 hours post treatment with UNSW-SC-22 at the indicated doses. **(C)** Cell cycle assay with propidium iodide indicating the proportion of cells in sub-G_1_ after 24 hours treatment with UNSW-SC-22 at the indicated doses. **(D)** Apoptosis assay with annexin-V/7-AAD staining in D283MED cells represented in stacked column graphs indicating proportion of apoptotic cells after treatment with UNSW-SC-22 for 48 hours at the indicated doses. **(E)** Immunoblot of PARP, cleaved-PARP, p21, caspase-3, and cleaved-caspase-3 proteins in the indicated treatment groups for D283MED and CHLA-01-MED cells. **(F)** Immunoblot H2AX and γH2AX proteins in the indicated treatment groups for D283MED cells. Reported p-values are from one-way ANOVA, error bars represent the standard error of the mean (SEM) of at least three independent biological repeats. (*\*p<0.1, **p<0.01, ***p<0.001, **** p< 0.0001*). The loading controls in Figure 4F were reused here in Figure 5F as samples were run on the same gel and are shown in separate panels for clarity.

### UNSW-SC-22 Combination Therapy with Histone Deacetylase Inhibitors Entinostat and Quisinostat in MYC-Driven Medulloblastoma Cell Lines

We have shown UNSW-SC-22 to be effective *in vitro as* a single agent in MYC-driven MB cells **(Fig.1, A and B)**. Since UNSW-SC-22 is an analogue of SE486-11, which synergises with HDAC inhibitors ^17,18^, We next determined whether UNSW-SC-22 could enhance the effects of two HDAC inhibitors: the class I HDAC inhibitor entinostat and the pan-HDAC inhibitor quisinostat. A panel of MB cell lines was first treated with single-agent entinostat (0-20 μM) or quisinostat (0-0.25 μM) for 72 hours, resulting in dose-dependent cytotoxicity across all tested MB cell lines (**Fig. S6A**). Similar treatment of normal fibroblast and human astrocyte cell lines showed a favourable therapeutic window (**Fig. S6B**). We then used a 5 × 6 viability matrix, with concentrations flanking the IC₅₀ of each drug, to identify dose ratios with the greatest synergy across MB cell lines. For each drug pair, a synergy index was calculated using two models: excess over highest single agent and Bliss independence. A synergy index >0 indicated synergy, =0 indicated additivity, and <0 indicated antagonism (**Table 3 and 4, Fig. S6C to F**). UNSW-SC-22 synergistically enhanced the cytotoxicity of both entinostat and quisinostat across MB cell lines at different doses. A theoretical Bliss viability line was generated for each combination, with values below the line indicating synergy (**Fig. S6G and H**). To further validate this synergy, each combination was re-tested using an expanded nine-point dose range based on the identified synergistic dose ratio. This confirmed clear synergy between UNSW-SC-22 and entinostat, and between UNSW-SC-22 and quisinostat, in MYC-driven MB cell lines (**Fig. 6A and B, Fig. S6I and J**). In contrast, the same dose ratios caused only a ∼30% reduction in viability in normal MRC5 fibroblasts (**Fig. 6C and D**). Combination treatment with entinostat, but not quisinostat, produced the greatest reduction in MYC protein levels and the strongest induction of DNA damage and apoptosis (**Fig. 6E to H, Fig. S6K to N**). This is consistent with previous reports that class I HDACs are frequently highly expressed in MYC-driven MB ^32,33^. This may explain the stronger synergy observed with entinostat compared with quisinostat. Together, our findings suggest that combining UNSW-SC-22 with entinostat may represent a promising therapeutic strategy for MYC-driven MB.

**Figure 6.**
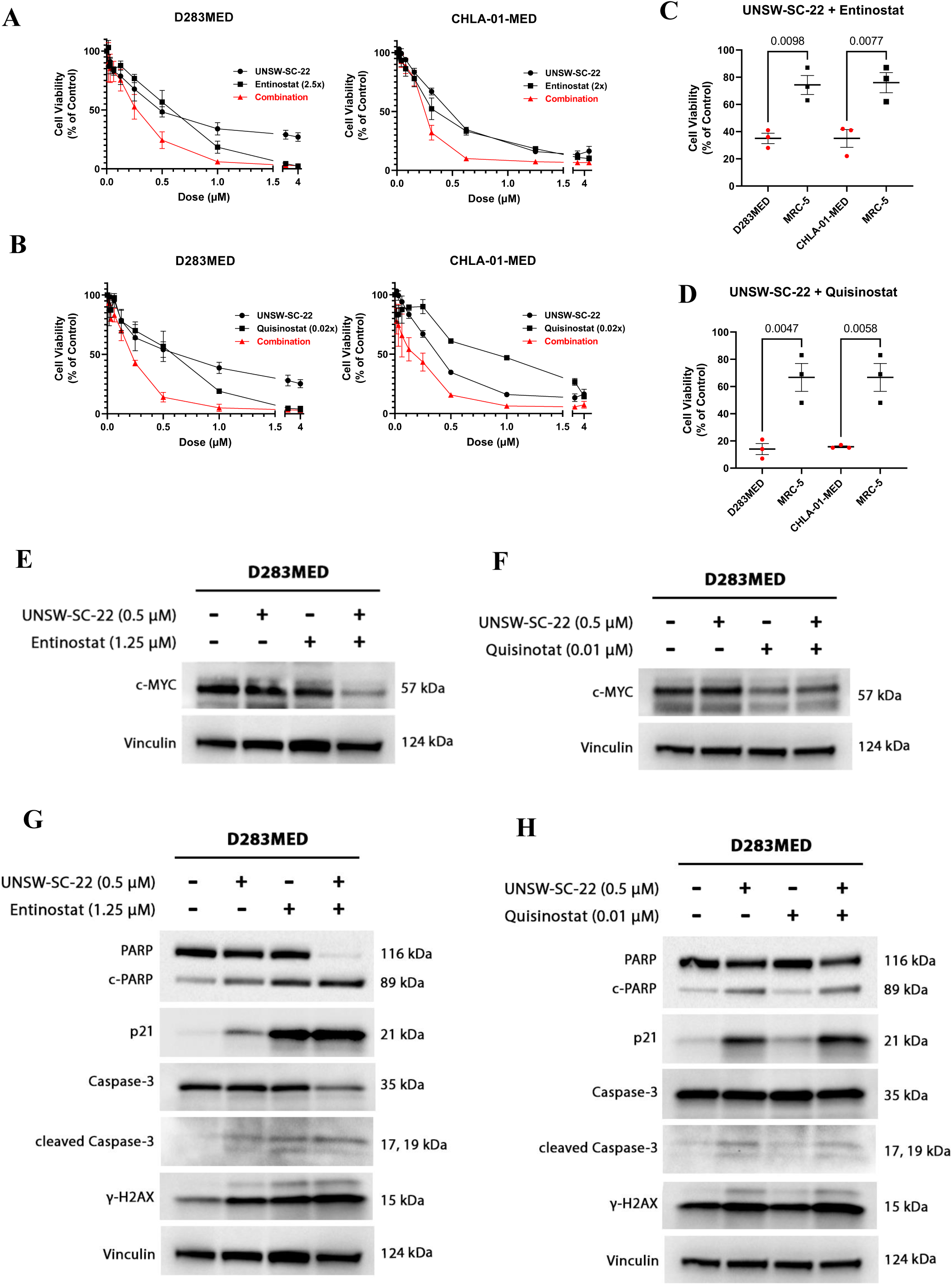
UNSW-SC-22 synergises with histone deacetylase inhibitors entinostat and quisinostat in MYC-driven medulloblastoma cell lines. **(A-B)** Cell viability curves of D283MED and CHLA-01-MED MB cell lines upon treatment with UNSW-SC-22 and **(A)** entinostat or **(B)** quisinostat for 72 hours in their determined synergistic ratio respectively. Values represent cell viability relative to DMSO vehicle control. **(C-D)** Comparison of cell viability after treatment with a combination of UNSW-SC-22 and **(C)** entinostat or **(D)** quisinostat at their determined synergistic dose ratio in MB cell lines and 1 FB cell line (MRC-5). **(E-F)** Immunoblot of MYC proteins for D283MED and CHLA-01-MED cells after 72-hours treatment with UNSW-SC-22 and **(E)** entinostat or **(F)** quisinostat at the indicated doses. **(G-H).** Immunoblot of PARP, cleaved-PARP, p21, caspase-3, cleaved-caspase-3 and γH2AX proteins for D283MED and CHLA0-01-MED cells after 24-hours treatment with UNSW-SC-22 and **(G)** entinostat or **(H)** quisinostat at the indicated doses. The curves represent non-linear regression fits of normalized cell viability data derived from three individual runs. Error bars represent the standard error of the mean and statistical comparisons were performed using one-way ANOVA. The loading controls in Figure 6E and 6F were reused in Figure 6G and 6H as samples were run on the same gel and are shown in separate panels for clarity.

### UNSW-SC-22 Crosses the BBB And Reduces Tumorigenicity in Vivo

We next evaluated the *in vivo* efficacy of UNSW-SC-22 in MYC-driven MB models. Pharmacokinetic analysis in C57BL/6 mice showed effective BBB penetration following intraperitoneal dosing at 60 mg/kg. UNSW-SC-22 reached peak plasma and brain concentrations of 8.46 μM and 10.38 μM, respectively, at 15 minutes, with a peak brain-to-plasma ratio of 1.44 (**Fig. 7A and B**). Brain exposure was briefly maintained after plasma levels declined, supporting its potential for treating central nervous system malignancies. We then assessed UNSW-SC-22 in an orthotopic MYC-amplified MB xenograft model. Balb/c nude mice were injected with 50,000 D425MED-Luc cells, resulting in 100% tumour engraftment, although with high inter-mouse variability (**Fig. S7A**). Mice were treated with DMSO or UNSW-SC-22 at 90 mg/kg intraperitoneally on a 5-days-on, 2-days-off schedule for 21 days. UNSW-SC-22 increased median overall survival by 2 days, although this was not statistically significant (**Fig. S7B**). However, treatment significantly improved event-free survival, defined by the onset of mild neurological symptoms such as head tilt, ataxia or circling (p = 0.0335; **Fig. 7C**). UNSW-SC-22 also reduced bioluminescence during treatment compared with DMSO (**Fig. 7D, Fig. S7C**), with significantly lower tumour signal observed on day 8 (p = 0.0079; **Fig. 7E**). These findings suggest that UNSW-SC-22 penetrates the BBB and can suppress MYC-driven MB tumour progression, with potential for improved efficacy through extended dosing or combination therapy. Immunohistochemical analysis of endpoint brain sections was performed to assess apoptosis and proliferation following treatment with DMSO or UNSW-SC-22 at 90 mg/kg/day. CC3 and MYC levels were unchanged, while Ki67 expression was reduced in UNSW-SC-22-treated mice (**Fig. S7D**). H&E staining also showed reduced brain haemorrhage and fewer mitotic cells following UNSW-SC-22 treatment (**Fig. S7E**).

**Figure 7.**
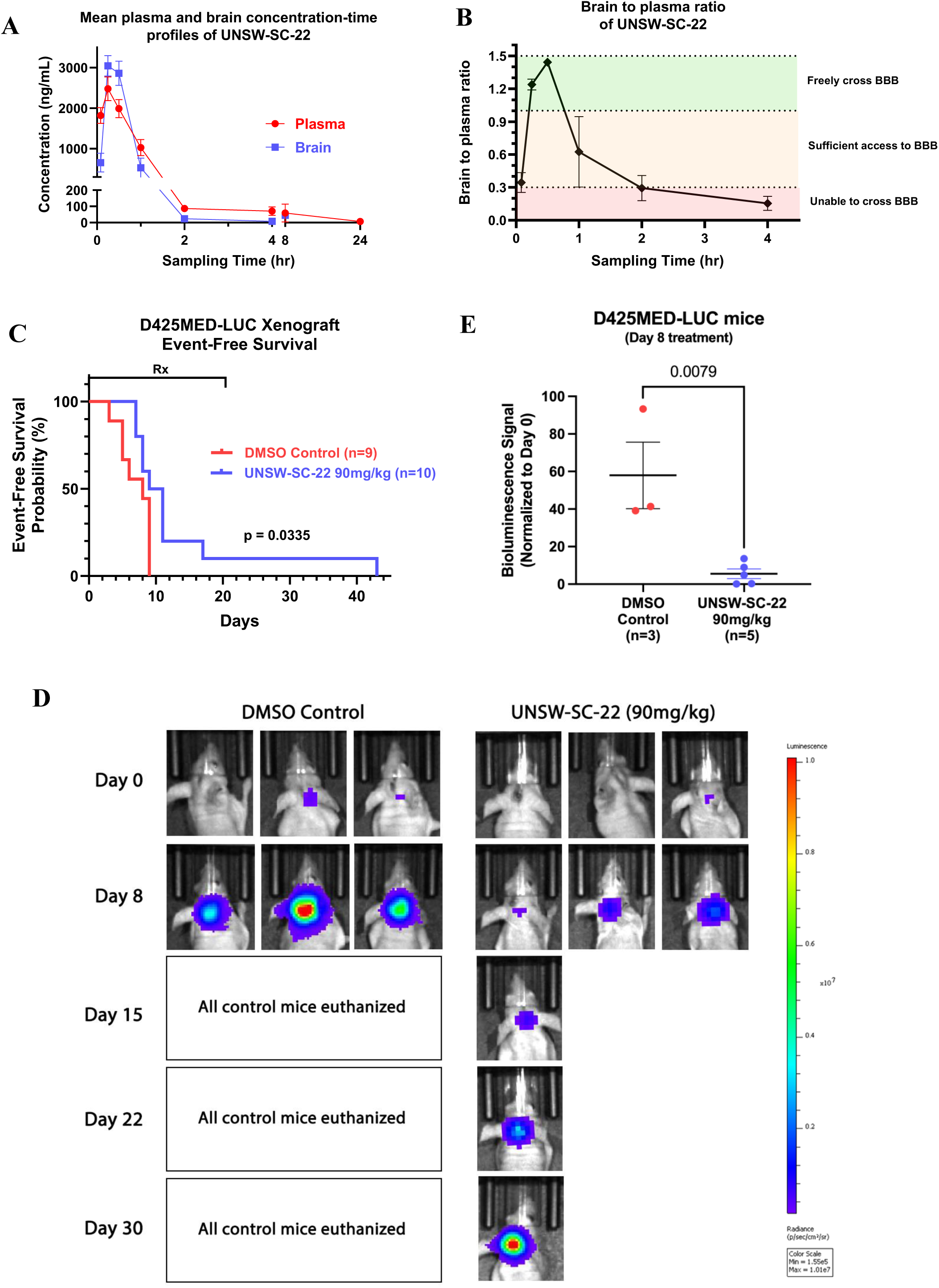
UNSW-SC-22 crosses the blood brain barrier and slows down tumour growth in D425MED orthotopic xenograft mice. **(A)** Mean plasma and brain concentration-time profiles of UNSW-SC-22 at 0.083, 0.25, 0.5, 1, 2, 4, 8 and 24 hours after intraperitoneal administration of a single dose of 60mg/kg in male C57BL/6 mice. **(B)** The brain to plasma ratio of UNSW-SC-22 across different sampling times. Error bars represent the standard error of the mean for triplicate runs. **(C)** Kaplan-Meier event-free survival curves of Balb/c nude mice xenografted with D425-LUC cells, post tumour formation during the administration of DMSO control or single agent UNSW-SC-22 (90mg/kg IP) for 21 days on 5 days on, 2 days off treatment schedule. **(D)** Representative bioluminescence images of D425-Luc orthotopic xenograft mice during treatment with DMSO control or UNSW-SC-22 (90mg/kg IP). **(E)** Comparison of normalized bioluminescence signal for each mouse during treatment with DMSO control (n=3) or UNSW-SC-22 90mg/kg (n=5). Error bars represent the standard error of the mean for at least three independent mice. P-value is a result of comparisons from unpaired t-tests.

The MYC orthologue, MYCN, is often amplified in high-risk or relapsed MB, particularly in the sonic hedgehog (SHH) and Group 4 subtypes ^34,35^. Hence, we determined whether UNSW-SC-22 would be effective in MYCN-driven MB. GTML/p53ER mice are a cross between mice homozygous for a doxycycline-regulable MYCN transgene driven by the Glt1 tissue-specific promoter (GTML; *Glt1-tTA/TRE-MYCN-LUC*) ^23^, and a p53 mutant mouse model (p53ER; *Trp53^KI/KI^*), where the endogenous *Trp53* gene is replaced with an inducible p53 knock-in allele (*Trp53^KI^*) encoding a 4-hydroxytamoxifen (4-OHT)-regulable mutant p53ER^TAM^ fusion protein ^24,36^. The homozygous (GTML/*Trp53^KI/KI^*) mice had a median survival time of 52 days and a significant decrease in overall survival (p < 0.0001) in comparison to heterozygous and wild type mice **(Fig. 8A)**, consistent with previous reports ^37^. Immunohistochemical staining of brain tumours from homozygous mice showed tumour-specific MYCN expression (**Fig. S8A**). We next assessed UNSW-SC-22 in GTML/Trp53^KI/KI^ MB-derived neurospheres, in which MYCN expression can be fully repressed by doxycycline (**Fig. 8B, Fig. S8B and C**). Neurospheres were treated with DMSO or doxycycline for 72 hours to repress MYCN, followed by UNSW-SC-22 treatment at 0-20 μM for another 72 hours. UNSW-SC-22 showed >30-fold greater growth inhibition in MYCN-expressing neurospheres compared with MYCN-repressed neurospheres (**Fig. 8C, Fig. S8D**). These findings further confirm that UNSW-SC-22 activity is MYCN-dependent and remains cytotoxic in the absence of functional p53 signalling.

**Figure 8.**
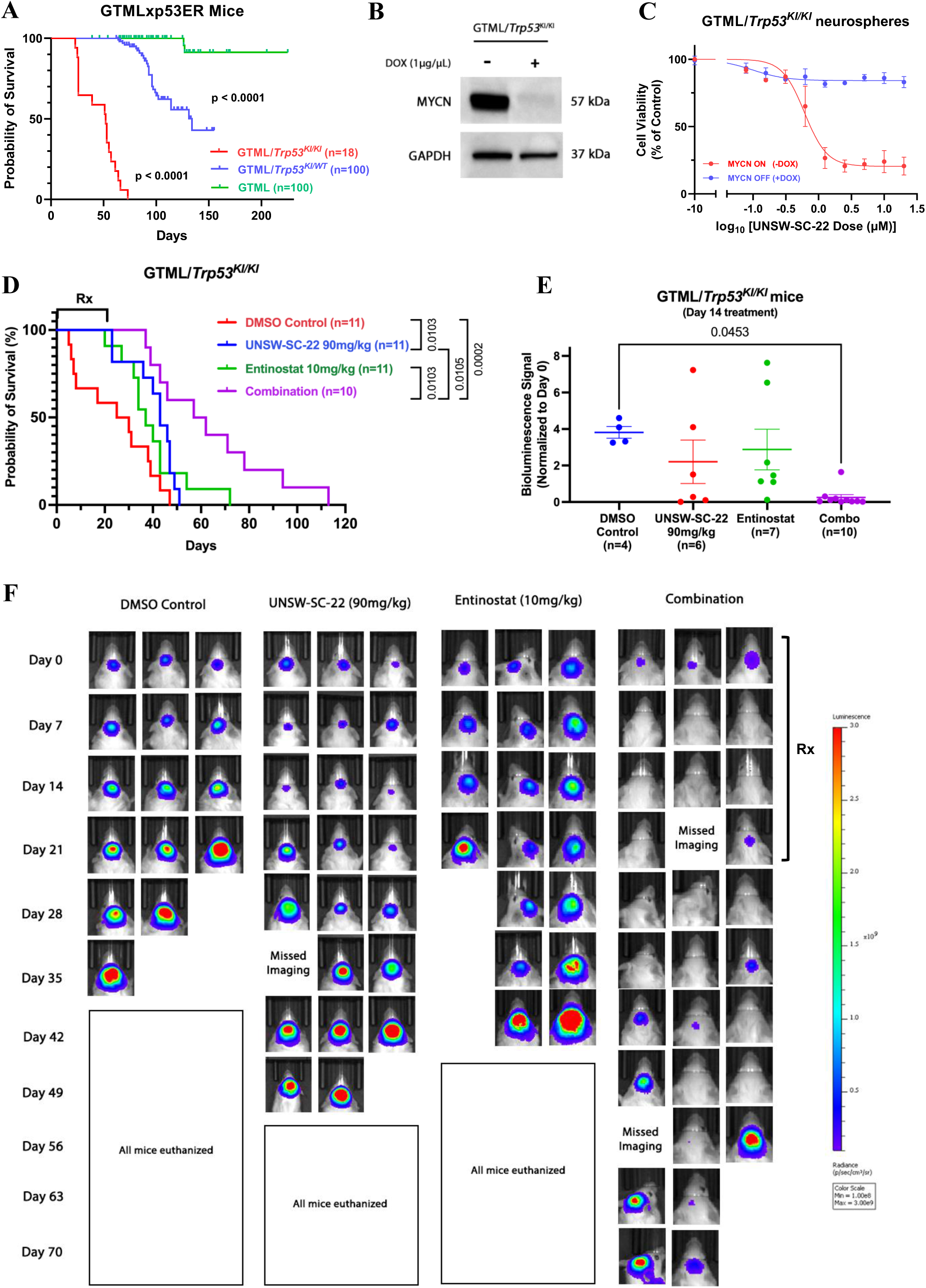
UNSW-SC-22 as a single agent and in combination with entinostat reduces MYCN-driven medulloblastoma tumour growth and prolongs survival in mice. **(A)** Kaplan-Meier survival curves for GTML/*Trp53KI/KI* (n=18), GTML/*Trp53KI/WT* (n=100) or GTML transgenic mice (n=100) as indicated. **(B)** Immunoblot of MYCN protein for GTML/*Trp53KI/KI* derived neurospheres cells in the presence or absence of 1μg/mL of doxycycline for 72 hours. **(C)** Cell viability curves of GTML/*Trp53KI/KI* neurospheres upon treatment with UNSW-SC-22 (0 – 20 μM) for 72 hours in the presence or absence of 1μg/mL of doxycycline as indicated. Values represent cell viability relative to a DMSO vehicle control. **(D)** Kaplan-Meier survival curves of GTML/*Trp53KI/KI* mice, during the administration of DMSO control, single agent UNSW-SC-22 (90mg/kg IP), single agent entinostat (10mg/kg IP) or a combination of UNSW-SC-22 and entinostat for 21 days of 5 days on, 2 days off treatment schedule. **(E)** Comparison of normalized bioluminescence signal for each mouse during day 14 of treatment with DMSO control (n=4), UNSW-SC-22 90mg/kg (n=6), entinostat 10mg/kg (n=7), or combination (n=10). **(F)** Representative bioluminescence images of GTML/*Trp53KI/KI* mice during treatment with DMSO, 90mg/kg/day UNSW-SC-22, 10mg/kg/day entinostat or a combination for a period of 21 days. Error bars represent the standard error of the mean for at least three independent mice. P-value is a result of comparisons from unpaired t-tests.

GTML/*Trp53^KI/KI^* mice were then treated with either DMSO, UNSW-SC-22 (90 mg/kg), entinostat (10 mg/kg) or a combination of UNSW-SC-22 and entinostat by intraperitoneal injection on a schedule of 5 days on, 2 days off, for a total of 21 days. UNSW-SC-22 alone and in combination with entinostat significantly prolonged survival in GTML/*Trp53^KI/KI^* compared to DMSO vehicle control (p = 0.0103, p = 0.0002), **(Fig. 8D).** When comparing bioluminescence signals between treatment groups on day 14 of treatment, mice treated with the combination had significantly lower signals compared to DMSO vehicle control (p = 0.0453) **(Fig. 8E)**. Although, we observed a reduction in tumour bioluminescence in GTML/Trp53^KI/KI^ mice during the treatment period, bioluminescence began increasing 1-2 weeks after treatment completion. **(Fig. 8F).** There were no signs of drug toxicity, indicating that UNSW-SC-22 alone and in combination with entinostat was well-tolerated **(Fig. S8E).** We observed no difference in the levels of CC3 or MYCN expression in UNSW-SC-22 treated mice compared to DMSO treated mice **(Fig. S7F)**.

## DISCUSSION

Here, we report a novel small-molecule MYC inhibitor that directly binds MYC/MYCN, promotes proteasomal degradation and suppresses MYC-dependent tumour growth. Its ability to cross the blood-brain barrier resulted in significant anticancer activity in murine models of MYC- and MYCN-driven medulloblastoma. These findings support the development of MYC-targeted degradation as a promising therapeutic strategy for high-risk medulloblastoma and other MYC/MYCN-driven cancers.

We showed that UNSW-SC-22 directly binds MYC, with target engagement further validated in medulloblastoma cells using DARTS. Although its binding affinity was lower than some previously reported MYC inhibitors, UNSW-SC-22 significantly reduced MYC protein levels and showed clear MYC-dependent activity in medulloblastoma cells. While MYC mRNA was not transcriptionally suppressed, MYC target genes were downregulated, consistent with reduced MYC protein stability. This was associated with increased MYC Thr58 phosphorylation following UNSW-SC-22 treatment. We also identified USP1 as a downstream transcriptional target repressed by UNSW-SC-22. Given that USP1 can stabilise the MYC/MAX complex by reducing ubiquitination, its suppression may further contribute to MYC destabilisation^28^, so UNSW-SC-22 based degradation of MYC could propagate USP1 depletion and a feedback loop that reinforces MYC degradation signals. MYC promotes cell cycle progression through the repression of cell cycle inhibitors such as p21 and p27 at the G1/S cell cycle boundary ^38,39^. Our data highlights the prominent anti-proliferative effects of UNSW-SC-22 through induction of p21, akin to other MYC inhibitors such as Omo-myc, which was shown to induce cell cycle arrest by activation of p21 ^40^.

Class I HDACs (HDAC1, HDAC2 and HDAC3) are frequently overexpressed in MYC-driven MB, and have been shown to interact directly with MYC, enhancing its protein stability through complex formation ^32^. Recent studies have shown that both the class I HDAC inhibitor entinostat, and the pan-HDAC inhibitor quisinostat, can reduce MYC protein levels in MB cells ^41,42^. Moreover, entinostat is also currently being investigated in combination with nivolumab for use in children with high-risk cancers in the INFORM2 clinical trial (NCT03938042) ^43^. Our data suggest potent synergy between UNSW-SC-22 and HDAC inhibitors, particularly entinostat. This combination synergistically reduced MYC expression, increased DNA damage and induced apoptosis. Although the mechanism underlying enhanced MYC suppression remains to be defined, it may reflect the combined inhibition of distinct pathways that converge on MYC regulation, leading to pronounced antitumour effects.

One of the most important factors to consider when developing inhibitors targeted at brain cancer driver genes is the ability of the compound to cross the BBB. This is often measured by its brain-to-plasma ratio, where compounds with a brain-to-plasma ratio of greater than 1 can freely cross the BBB ^44^. Temozolomide, the first-line treatment when combined with radiation for glioblastoma, has been shown to have good BBB penetration, with a brain-to-plasma ratio of 1.1 at 0.5 hours following intraperitoneal administration of 266.7 mg/kg ^45^. UNSW-SC-22 showed favourable brain penetration, with a brain-to-plasma ratio of 1.25 at 0.25 hours after 60 mg/kg, comparable to temozolomide. Consistent with this, UNSW-SC-22 delayed tumour growth and improved survival in MYC- and MYCN-driven medulloblastoma mouse models. However, in the aggressive MYC-driven xenograft model, the survival benefit was limited, likely due to rapid tumour progression and the short treatment window, with only one mouse completing the full treatment course. Further evaluation in additional patient-derived xenograft models will better define the therapeutic scope of UNSW-SC-22 against MYC-driven medulloblastoma. In the MYCN-driven GTML/p53ER model, combining UNSW-SC-22 with entinostat produced greater tumour suppression and survival benefit than UNSW-SC-22 alone. Overall, these findings support UNSW-SC-22 as a promising monotherapy and combination strategy for high-risk MYC/MYCN-driven medulloblastoma and potentially other MYC/MYCN-driven cancers.

## Supporting information

RNA sequencing data

## Author Contributions

Conceptualisation: SW, SG, QW, DC, BC, GM, MWP

Methodology: SW, SG, MT, DZ, MWP, NK. LC, TL, MM

Investigation: SW, SG, NC, LD, TLN, JS, NJ, MT

Visualisation: SW, SG, BC, GM

Funding acquisition: BC, GM, MWP

Project administration: SW, BC, GM, MWP

Supervision: QW, BC, GM, MWP, TL

Writing – original draft: SW

Writing – review and editing: SW, SG, BC, GM, TLN, MWP, LD

Manuscript approval – all authors

## Conflict of Interest Statement

Authors declare that they have no Conflict of interests.

## Funding

NHMRC Project Grant (APP1125171; G.M. Marshall, B.B. Cheung)

Cancer Institute NSW Program grant in 2022 (G.M. Marshall, RG211809)

NHMRC Synergy Grant in 2023 (G.M. Marshall, RG220538)

Cancer Council NSW and Box Rallies Foundation Project Grants (B.B. Cheung, RG21-08, B.B. Cheung, RG22-09 and B.B Cheung RG25-04).

Can Too Foundation in 2022 (B.B. Cheung, RG22-09)

Tour de Cure Annual Grants in 2023, 2024 and 2025 (B.B Cheung, RSP-268-FY2023 and RSP-178-2024, G.M. Marshall, RSP-385-2025)

Kids’ Cancer Project and Neuroblastoma Australia in 2023 (B.B. Cheung, RG23161)

NHMRC MRFF Grant in 2021 (G.M. Marshall, RG210235)

NHMRC Investigator Grant in 2020 (G. M. Marshall, RG1194264). NHMRC Investigator Grant (M.W. Parker, APP1194263).

The authors thank the Steven Walter Children’s Cancer Foundation, Hyundai Help for Kids Foundation, Katoomba Op shop, Perpetual IMPACT philanthropy funding and Neuroblastoma Australia for their financial support. Funding from the Victorian State Government Operational Infrastructure Support Scheme to St. Vincent’s Instiute is acknowledged.

## Acknowledgements

We thank the members of the Embryonal Cancer and Therapy Prevention (ECTP) laboratory for discussion and assistance provided throughout this project. We thank Anick Standley, Lauren Grote and Thiago Arrivabene from the Children’s Cancer Institute Animal Facility for coordination and support throughout the animal studies. We thank UNSW Biological Resource Imaging Laboratory and UNSW Flow Cytometry Facility for assistance. This research made use of the ACRF Facility for Innovative Cancer Drug Discovery located at the Bio21 Institute. We acknowledge the Melbourne Protein Characterisation platform at Bio21 Institute for exceptional technical support.

## Ethics Statements

Studies were conducted under approved ACEC Approval and Animal Research Authority (ARA) and HREC protocols under Associate Professor Belamy Cheung and Professor Glenn Marshll at UNSW Sydney and The Sydney Children’s Hospital Network.

## Data and materials availability

All data associated with this study are present in the paper or the Supplementary Materials. Materials can be made available upon signing a material transfer agreement with standard provisions. Some restrictions may apply on the use of materials involving UNSW-SC compounds, except for research aimed at reproducing the findings reported in the paper.

## List of Supplementary Materials

Fig. S1 to S8

Data file S1

Table 1 to Table 4

Supplementary Methods and Materials

## SUPPLEMENTARY FIGURES

**Figure S1.**
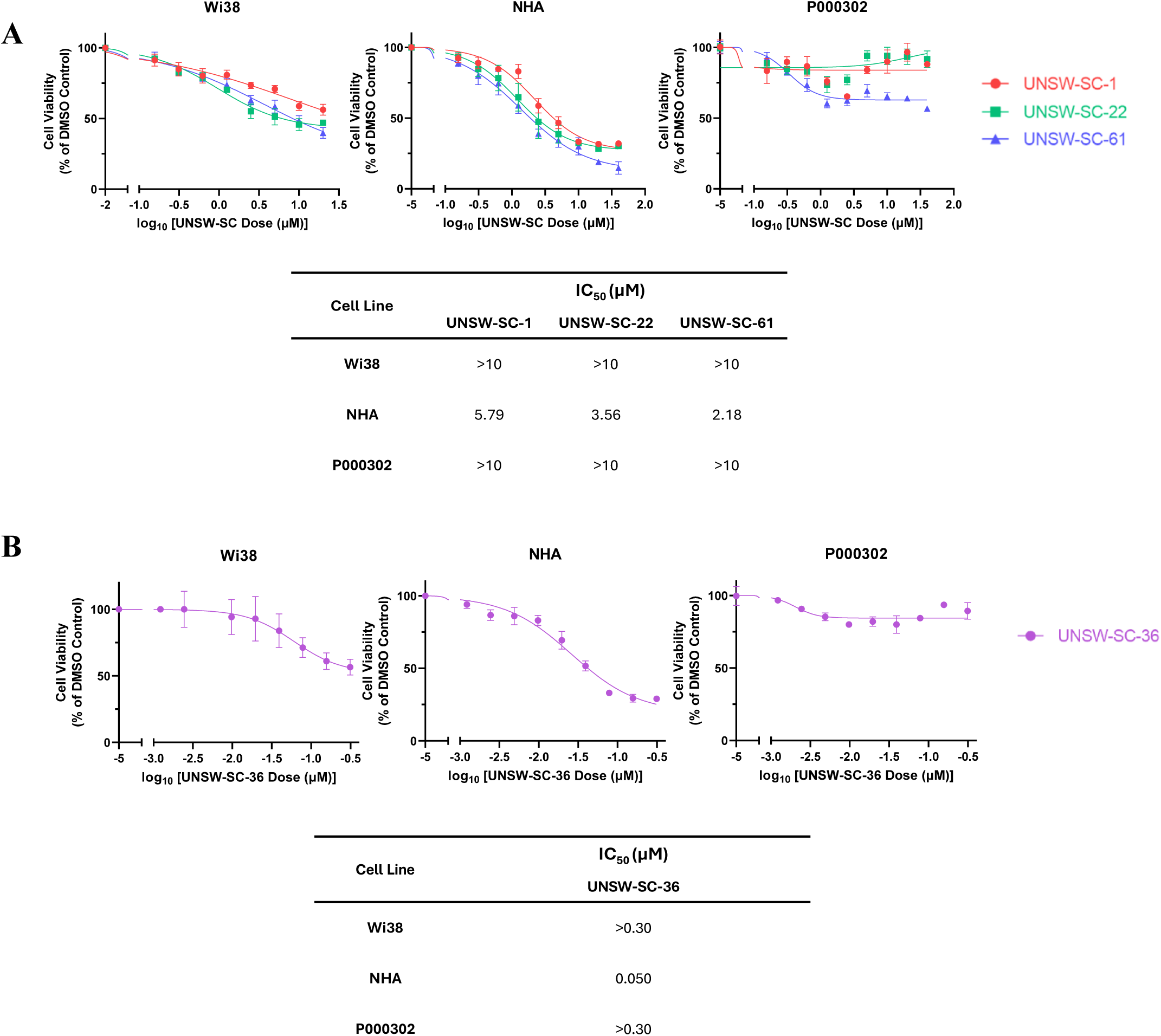
**(A)** Cell viability curves of human FB cell line Wi38 and 2 normal human astrocytes cell lines, NHA and P000302, upon treatment with UNSW-SC-1, UNSW-SC-22, and UNSW-SC-61 (0 – 20 μM) for 72 hours. Values represent cell viability relative to a DMSO vehicle control. **(B)** Cell viability curves of human FB cell line Wi38 and 2 normal human astrocytes, NHA and P000302 upon treatment with UNSW-SC-36 (0 – 0.3125 μM) for 72 hours and table of IC_50_ values.

**Figure S2.**
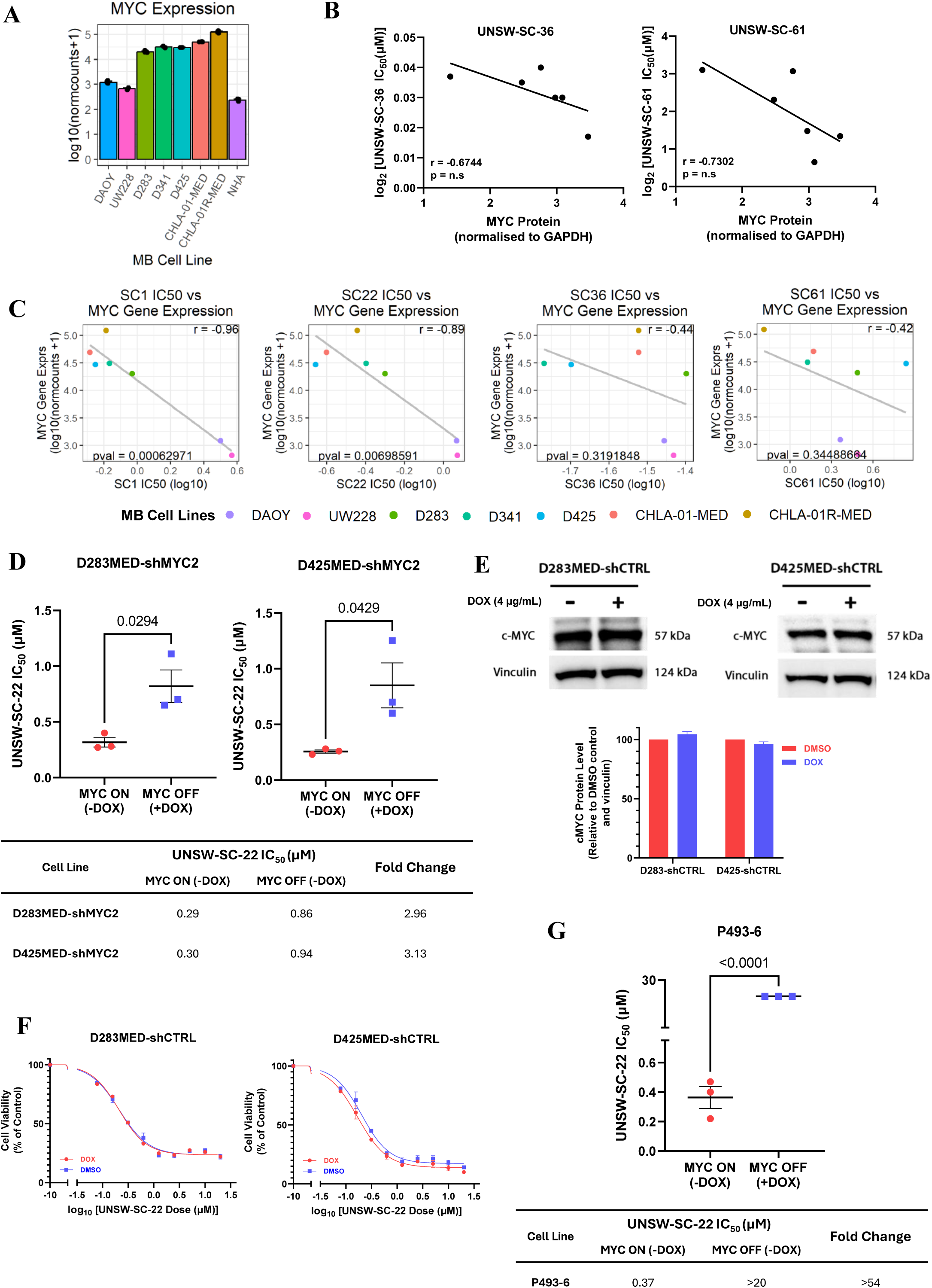
**(A)** The mRNA expression of MYC in 7 MB and 1 normal human astrocyte cell line analyzed from publicly available RNA-seq datasets (GSE185024, GSE171117, GSE52983, GSE102505). **(B)** Pearson correlation between MYC protein levels (from Figure 2A) and IC_50_ of UNSW-SC-36 and UNSW-SC-61. **(C)** Pearson correlation between MYC mRNA levels (from Figure S2A) in 7 MB cell lines and IC_50_ of all four UNSW-SC compounds. **(D)** IC_50_ comparisons for UNSW-SC-22 in shMYC MB cell lines in the presence or absence of MYC expression. **(E)** Immunoblot of MYC protein D283MED and D425MED shCTRL MB cell lysates in the presence or absence of 4μg/mL of doxycycline for 72 hours. (**F)** Cell viability curves of D283MED and D425MED shCTRL MB cell lines upon treatment with UNSW-SC-22 (0 – 20 μM) for 72 hours in the presence or absence of 4μg/mL of doxycycline. Values represent cell viability relative to a DMSO vehicle control. **(G)** IC_50_ comparisons for UNSW-SC-22 in P493-6 Burkitt’s lymphoma cells in the presence or absence of MYC expression. Each point represents average cell viability or IC50 values across at least three independent experiments. Error bars represent the standard error of the mean and statistical comparisons were performed using one-way ANOVA, two-way ANOVA, or t-test.

**Figure S3.**
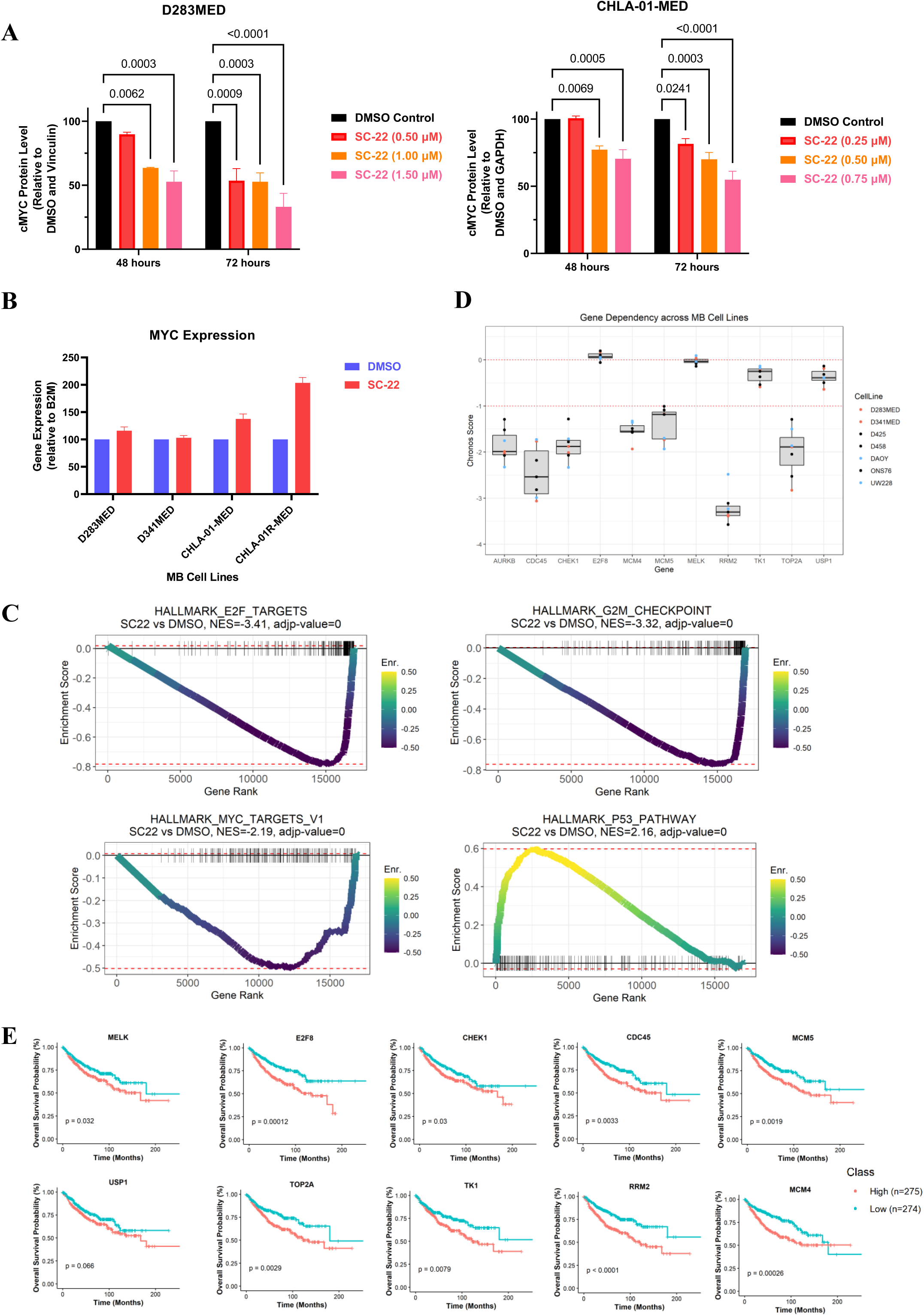
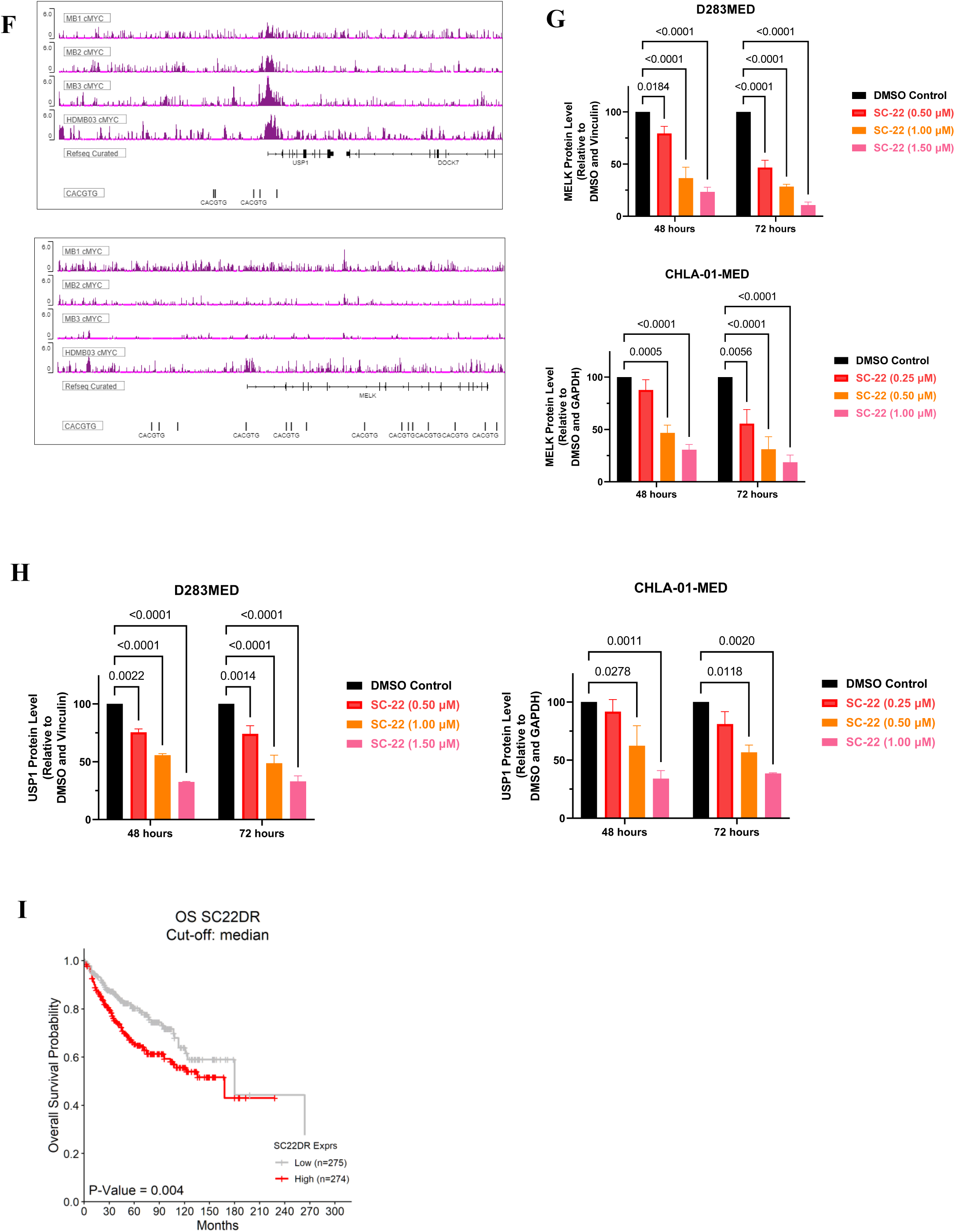
**(A)** Densitometry analysis of Figure 3A, with MYC proteins first normalized to vinculin, and then to their respective DMSO controls. **(B)** Real-time PCR analysis of mRNA expression for MYC following treatment of DMSO or 1.5 μM UNSW-SC-22 for 24 hours in high MYC expression MB cell lines D283MED, CHLA-01-MED, CHLA-01R-MED and D341MED cells. **(C)** Running enrichment plots across HALLMARK genesets; HALLMARK_E2F_TARGETS, HALLMARK_G2M_CHECKPOINT, HALLMARK_MYC_TARGETS and HALLMARK_P53_PATHWAY, with differentially expressed genes present in each gene set ranked by decreasing log2FC (UNSW-SC-22 vs DMSO control). The normalized enrichment score (NES) is indicated on the top of each plot. **(D)** The CRISPR dependency Chronos score for 10 candidate genes across 7 MB cell lines analyzed from the Cancer Dependency Map Portal (DepMap 24Q1). **(E)** Kaplan-Meier overall survival curves from Cavalli MB patient cohorts, split by a median gene mRNA expression cut-off for patient categorization of 10 candidate genes. **(F)** Analysis of publicly available ChIP-seq data (GSE143376) highlighting the canonical (CACGTG) MYC binding motifs located on the promoter sites of MELK and USP1 in 3 primary Group 3 samples (MB1, MB2, MB3) and MYC-amplified MB cell line (HDMB03). **(G-H)** Densitometry analysis of Figure 3F, with MELK and USP1 proteins first normalized to vinculin or GAPDH, and then to their respective DMSO controls. **(I)** Kaplan-Meier overall survival curves from Cavalli MB patient cohorts, split by a median gene mRNA signature (SC22DR) for patient categorization. Reported p-values are from one/two-way ANOVA, error bars represent the standard error of the mean (SEM) of at least three independent biological repeats.

**Figure S4.**
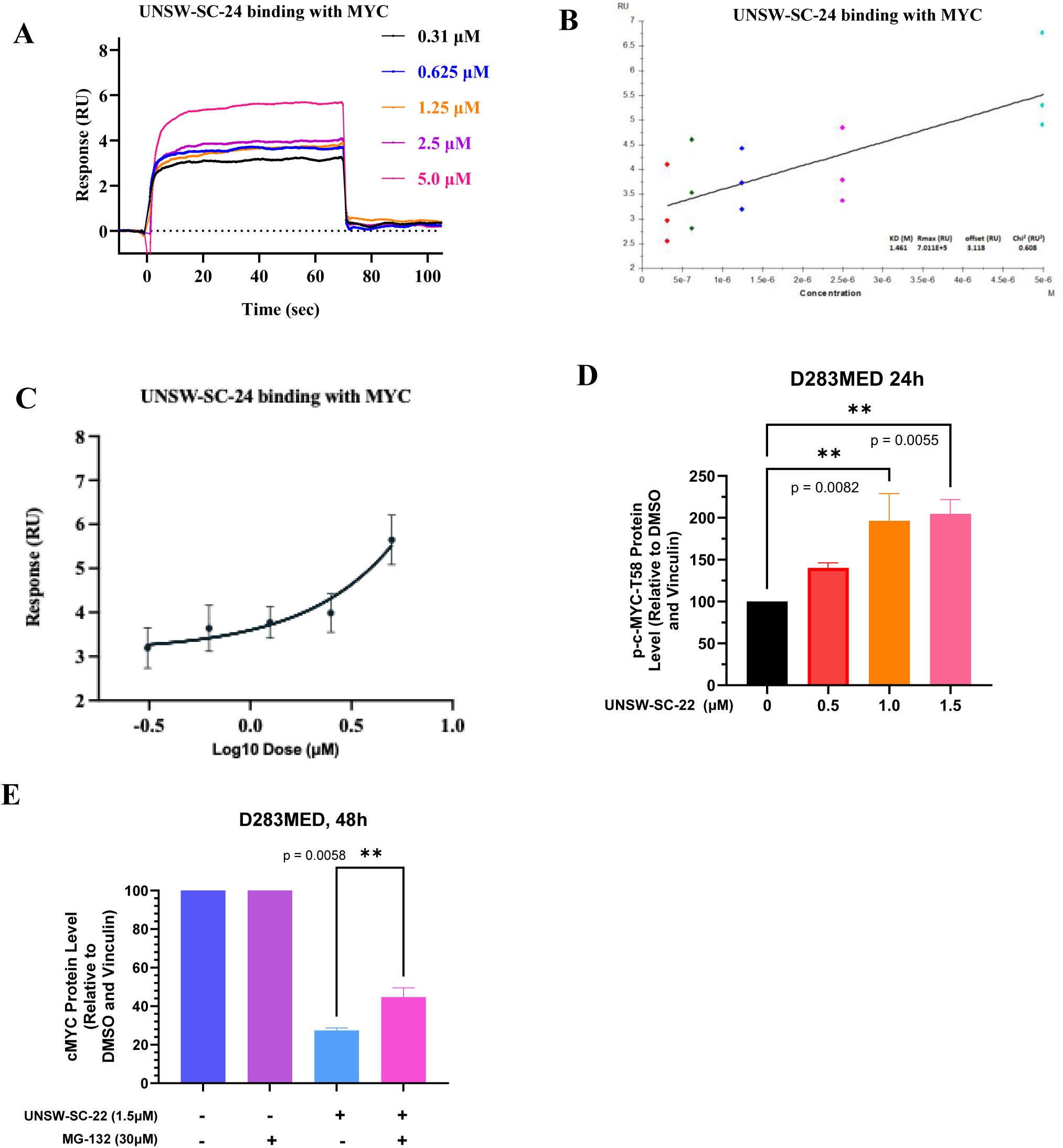
**(A-B)** Representative SPR curves illustrating the concentration-response binding of dead compound UNSW-SC-24 to MYC. **(C)** Normalized binding affinity curve showing the dose-response binding affinity between MYC and UNSW-SC-24. **(D)** Densitometry analysis of Figure 4G, where T58 MYC levels were first normalized to vinculin loading control, and then to DMSO control. **(E)** Densitometry analysis of Figure 4H, where MYC levels were first normalized to vinculin loading control, and then to their respective DMSO controls. Reported p-values are from one/two-way ANOVA, error bars represent the standard error of the mean (SEM) of at least three independent biological repeats.

**Figure S5.**
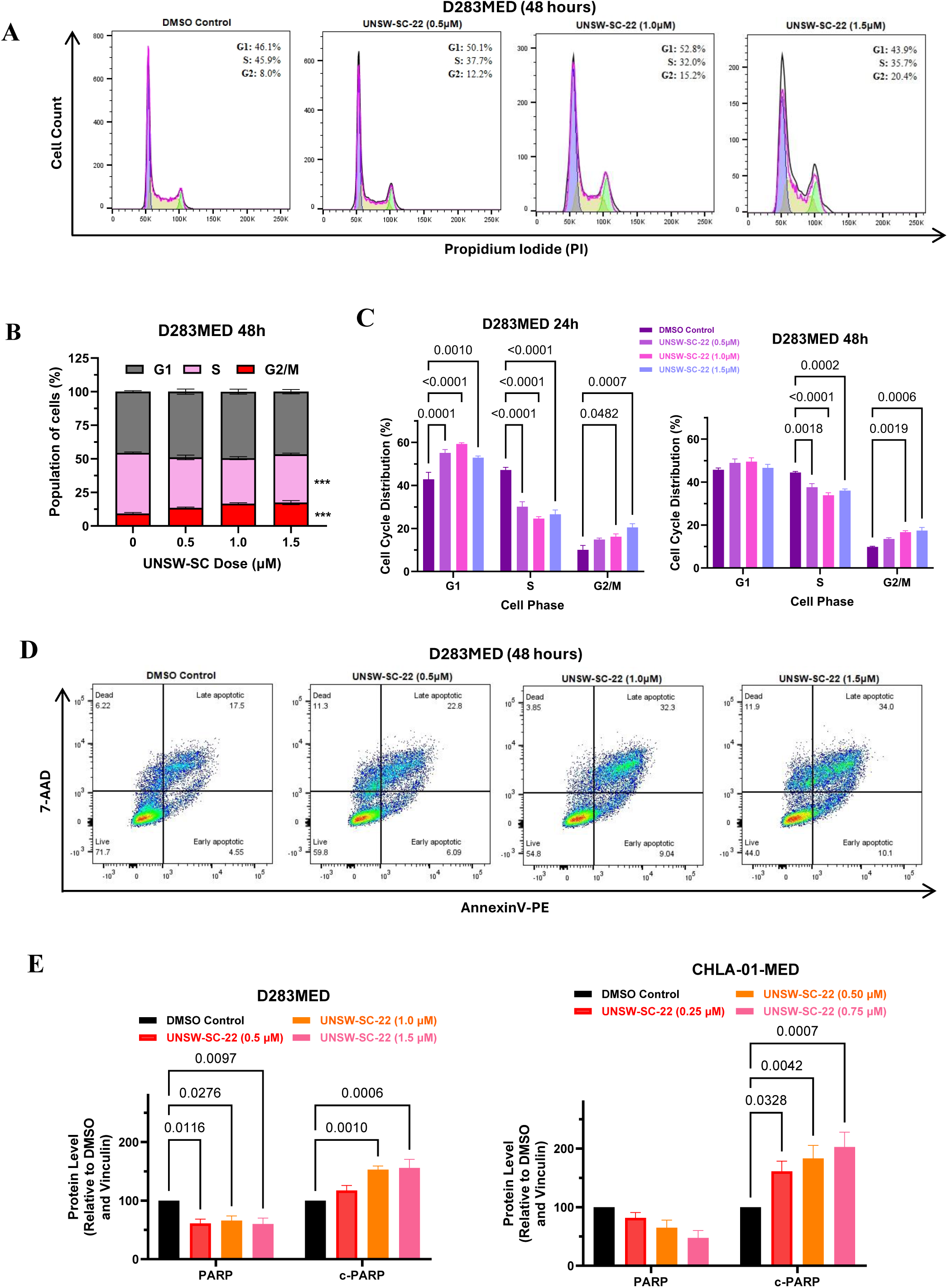
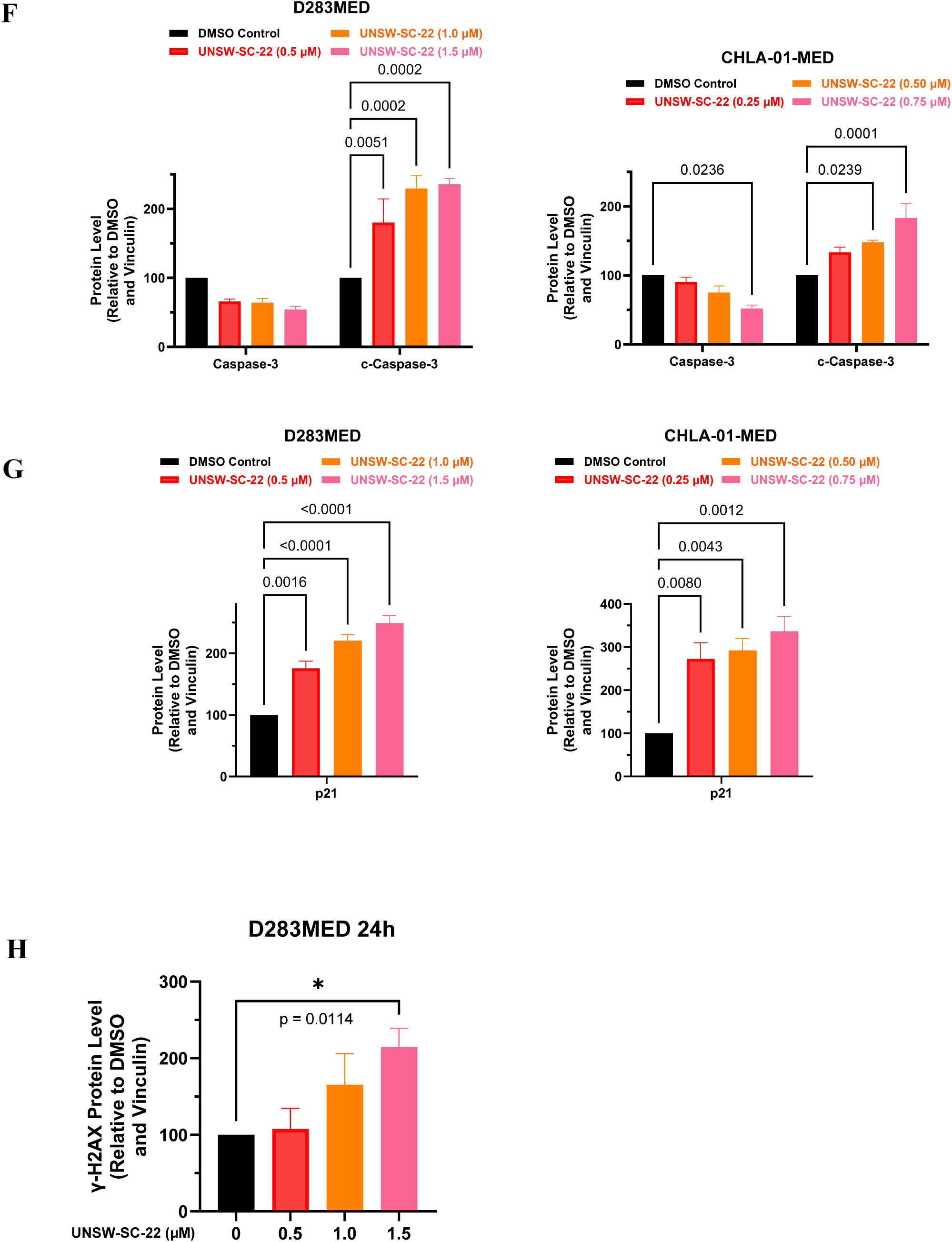
**A.** Flow cytometry analysis of cell cycle distribution in D283MED medulloblastoma cells after treatment with UNSW-SC-22 at the indicated doses for 48 hours, followed by propidium iodide staining. **B.** Cell cycle assay represented in stacked column graphs indicating the proportion of cells in each cell cycle phase 48 hours post treatment with UNSW-SC-22 at the indicated doses. **(C)** Cell cycle analyses representing the proportion of cells in each phase of the cell cycle post treatment with UNSW-SC-22 for 24 or 48 hours at the indicated doses in D283MED cells. **(D)** Representative quadrant gates for flow cytometry analysis of apoptosis with annexin-V/7-AAD staining in D283MED MB cells post treatment with UNSW-SC-22 for 48 hours at the indicated doses. **(E-G)** Densitometry analysis of Figure 5E, with PARP, cleaved-PARP, caspase-3, cleaved-caspase 3 and p21 proteins first normalized to vinculin, and then to their respective DMSO controls. **(H)** Densitometry analysis of Figure 5F, with γH2AX and H2AX proteins first normalized to vinculin, and then to their respective DMSO controls. Reported p-values are from one/two-way ANOVA, error bars represent the standard error of the mean (SEM) of at least three independent biological repeats.

**Figure S6.**
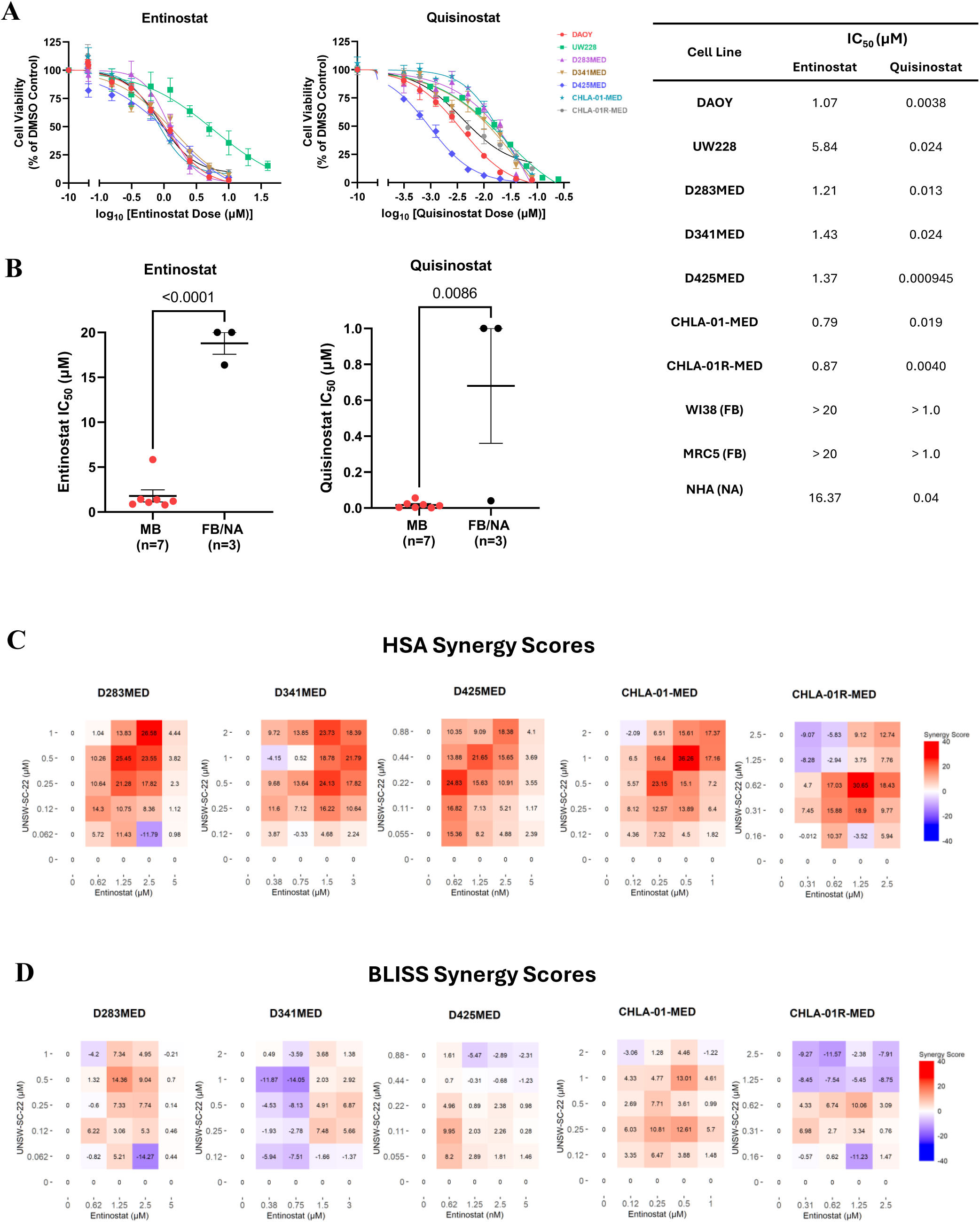
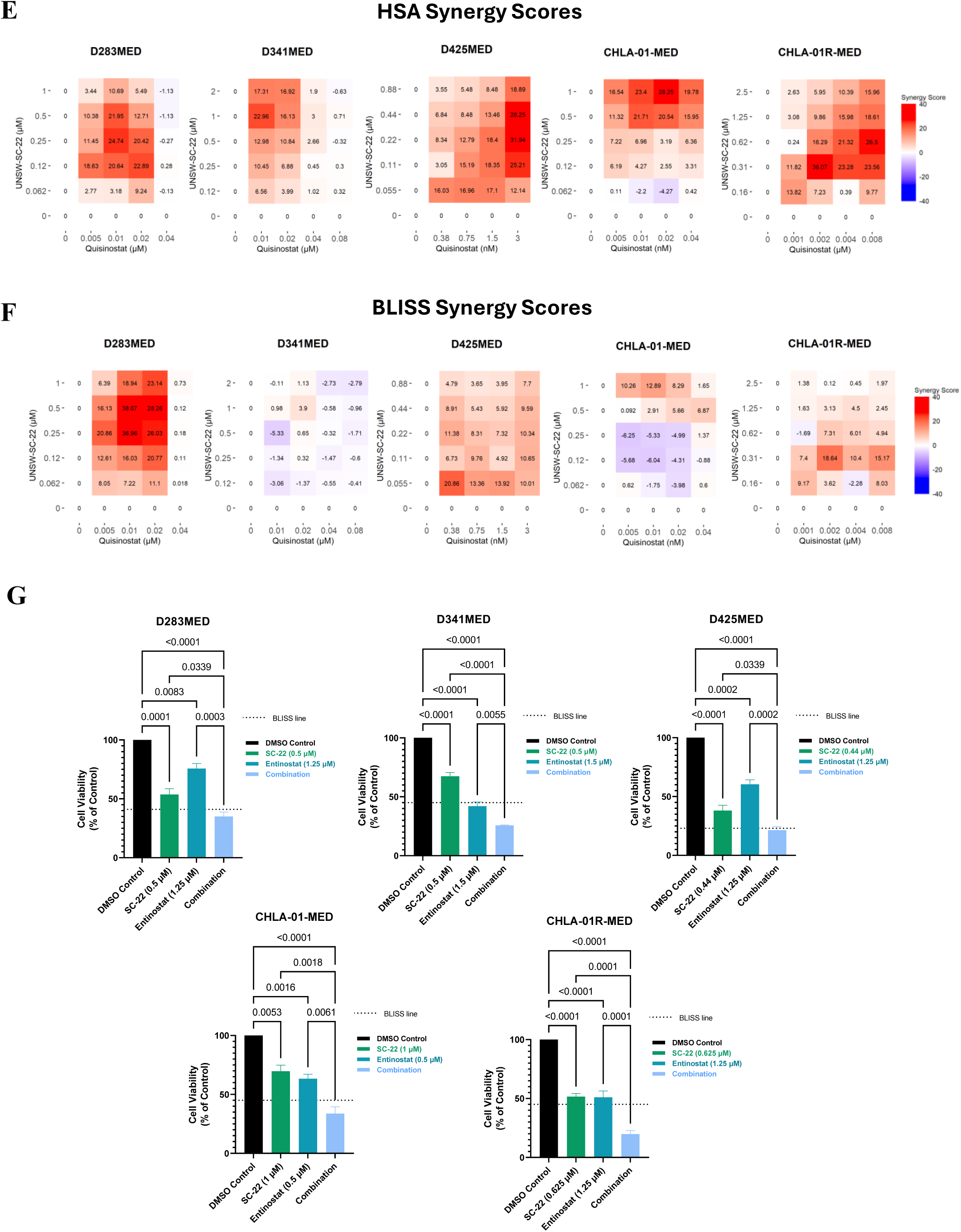
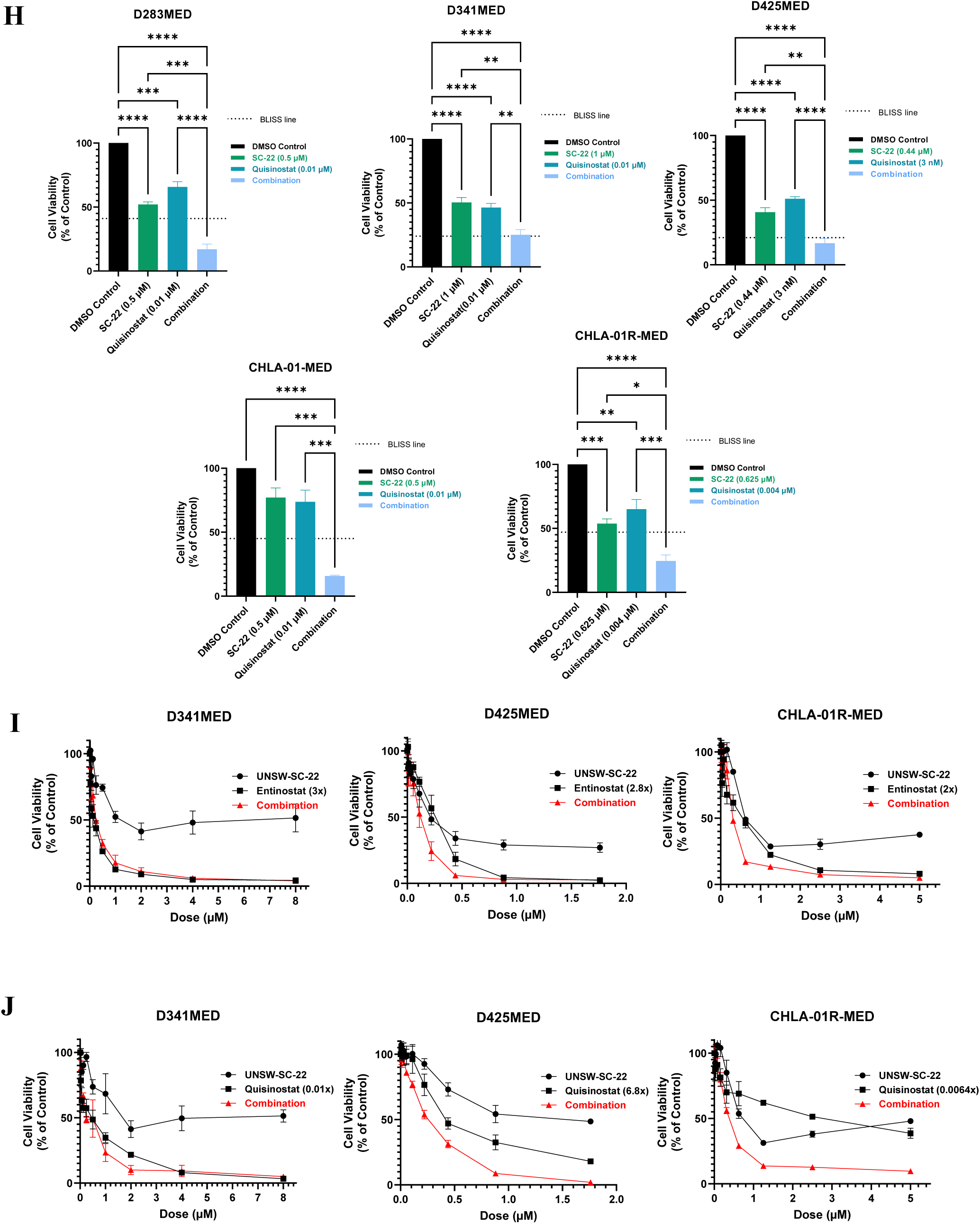
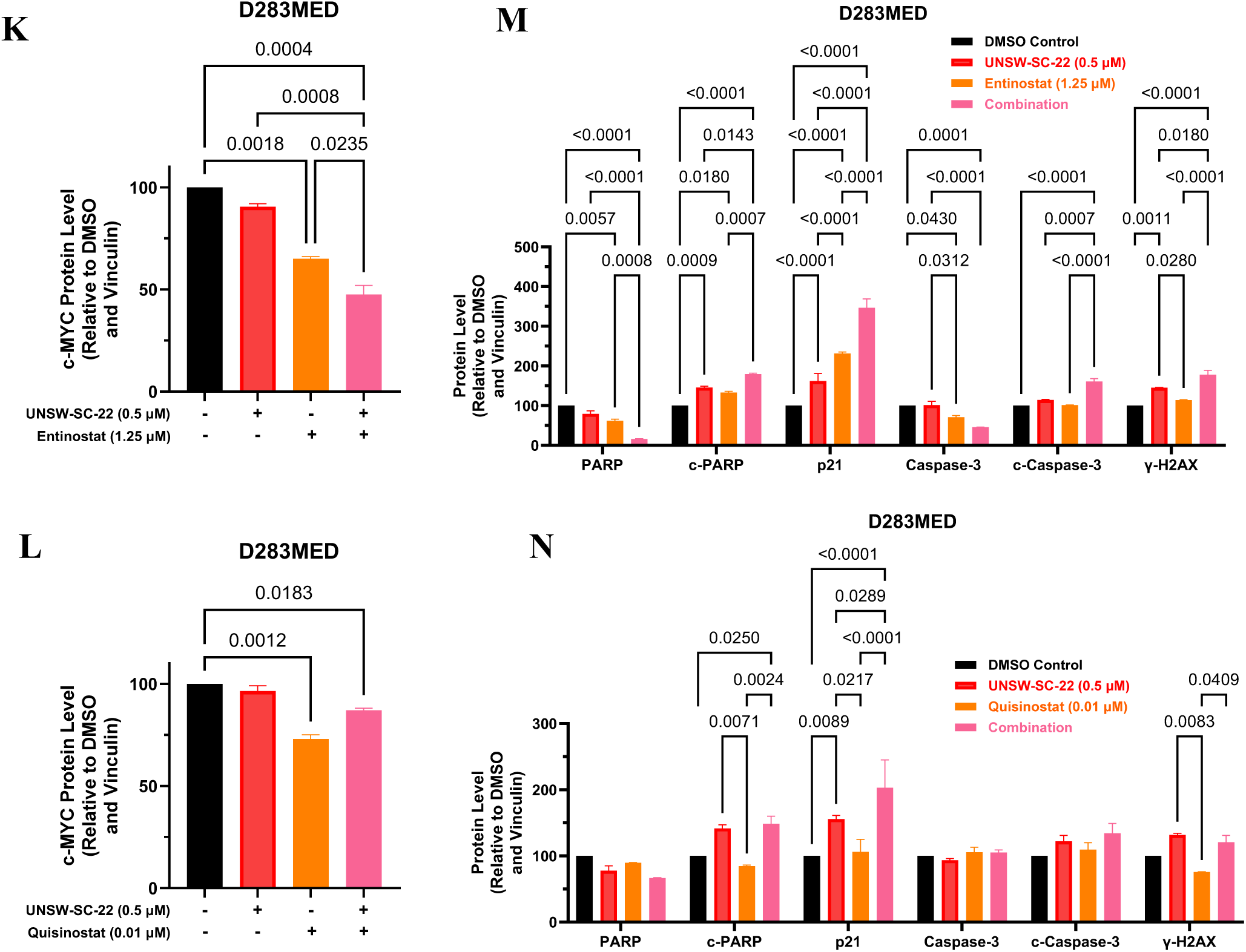
**(A)** Cell viability curves of 7 MB cell lines upon treatment with entinostat (0 – 20 μM) or quisinostat (0 – 0.25 μM) for 72 hours. Values represent cell viability relative to DMSO vehicle control. **(B)** Comparison of IC_50_ of entinostat or quisinostat in 7 MB cell lines, 2 fibroblast cell linea and 1 normal human astrocyte (FB/NA). Each point represents average cell viability or IC50 values across atleast three independent experiments. Error bars represent the standard error of the mean and statistical comparisons were performed using a t-test. **(C-F)** HSA and BLISS synergy scores calculated from a 6×5 matrix screen of UNSW-SC-22 and **(C-D)** entinostat or **(E-F)** quisinostat in 5 MYC-driven MB cell lines. For each plot, the synergy score provided is the synergism index (SI) expressed as a percentage (SI x 100). **(G-H)** Cell viability presented as a column graph at their determined synergistic dose ratio between UNSW-SC-22 and **(G)** entinostat or **(H)** quisinostat in 5 MYC-driven MB cell lines. For each plot, a “BLISS Line” is provided. **(I-J)** Cell viability curves of D341MED, D425MED and CHLA-01R-MED MB cell lines upon treatment with UNSW-SC-22 and **(I)** entinostat or **(J)** quisinostat for 72 hours in their determined synergistic ratio respectively. Values represent cell viability relative to DMSO vehicle control. **(K-L)** Densitometry analysis of **(K)** Figure 8E and **(L)** Figure 8F, with MYC first normalized to vinculin, and then to DMSO controls. **(M-N)** Densitometry analysis of **(M)** Figure 8G and **(N)** Figure 8H, with PARP, cleaved-PARP, caspase-3, cleaved-caspase 3, p21 and γH2AX proteins first normalized to vinculin, and then to their respective DMSO controls. Reported p-values are from one/two-way ANOVA, error bars represent the standard error of the mean (SEM) of at least three independent biological repeats.

**Figure S7.**
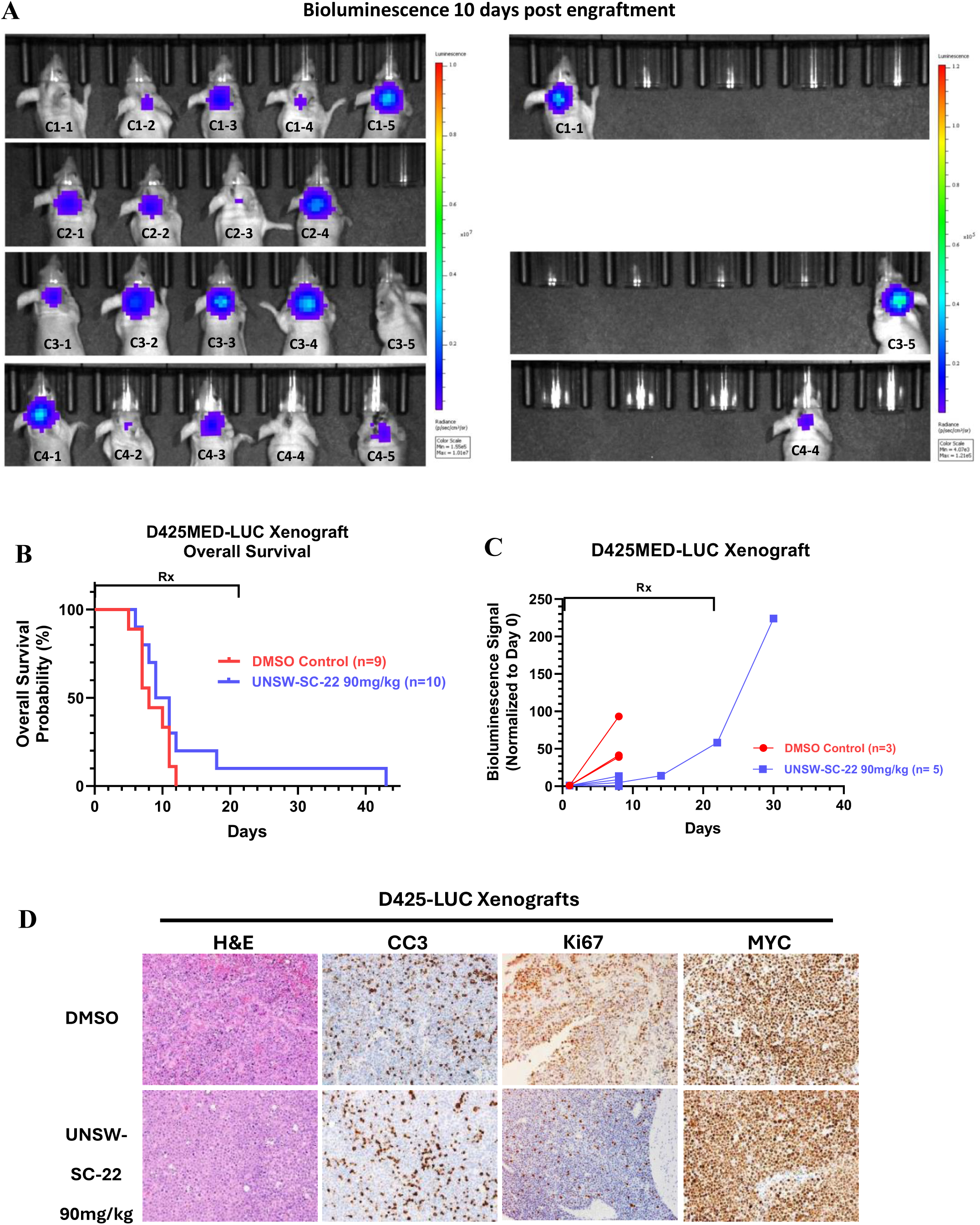
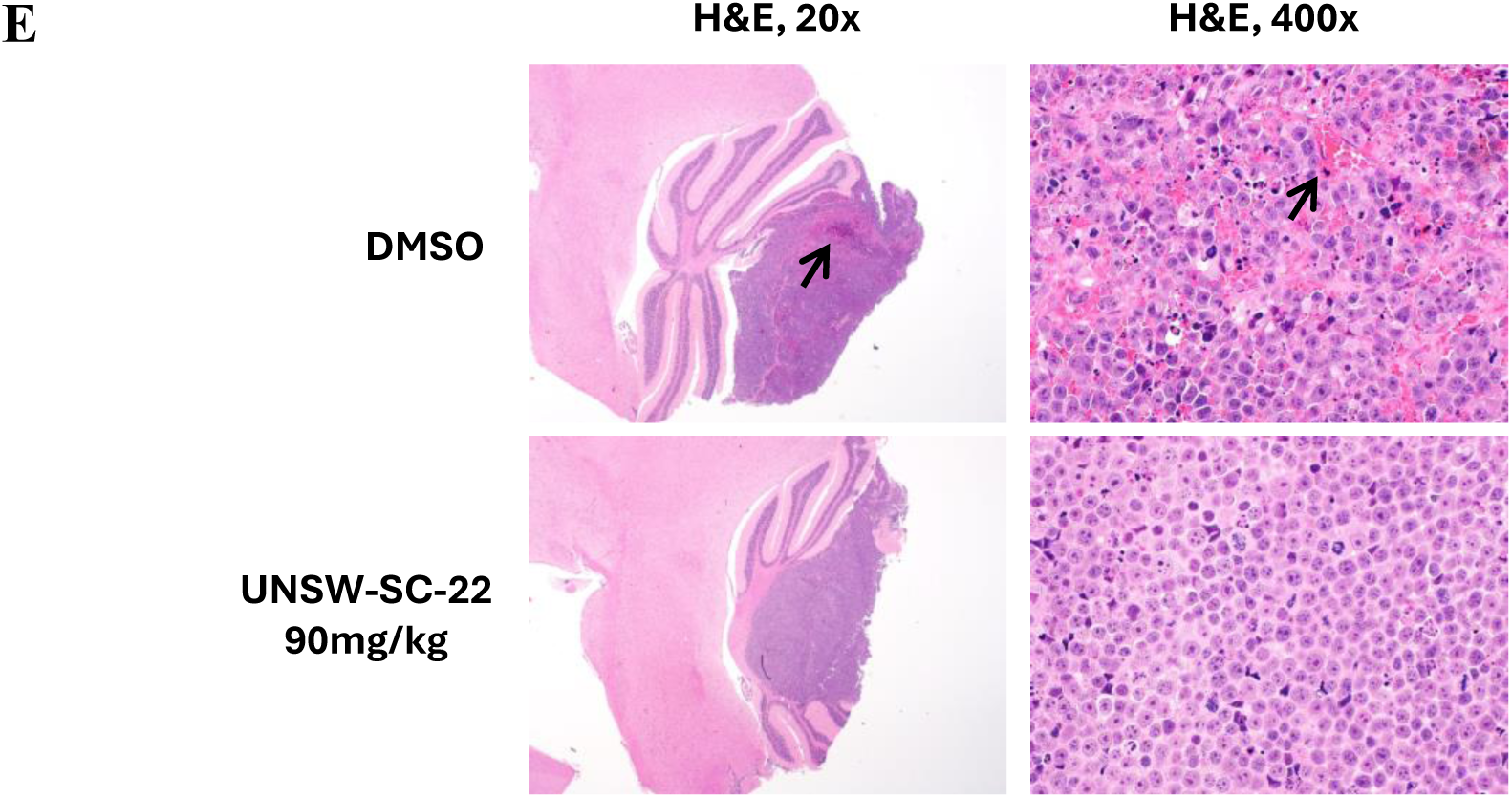
**(A)** Bioluminescence imaging of D425-Luc orthotopic xenograft mice 10 days after intracranial injection of D425-LUC MB cells without drug treatment. Individual mouse IDs are indicated by C1-1, C1-2, etc. Bioluminescence intensity is measured as radiance (p/sec/cm2/sr) represented by each scale bar as indicated. **(B)** Kaplan-Meier overall survival curves of Balb/c nude mice xenografted with D425-LUC cells, post tumour formation during the administration of DMSO control or single agent UNSW-SC-22 (90mg/kg IP) for 21 days on 5 days on, 2 days off treatment schedule. **(C)** Bioluminescence signals measured in radiance of D425MED-LUC orthotopic xenograft mice during treatment with DMSO control or UNSW-SC-22 (90mg/kg IP), normalized to day 0 of treatment for each mouse. **(D)** Representative images of UNSW-SC-22 treated D425-LUC xenograft brain sections after 21 days at magnification of 200x (20x objective lens, 10x eye piece lens) stained with H&E, anti-Cleaved caspase-3 (CC3), anti-Ki67 or anti-MYC. **(E)** Representative images of treated D425-LUC xenograft brain sections at magnification of 20x (2x objective lens, 10x eye piece lens) and 400x (40x objective lens, 10x eye piece lens) stained with H&E. Arrows indicate haemorrhage. Reported p-values are from one/two-way ANOVA, error bars represent the standard error of the mean (SEM) of at least three independent biological repeats.

**Figure S8.**
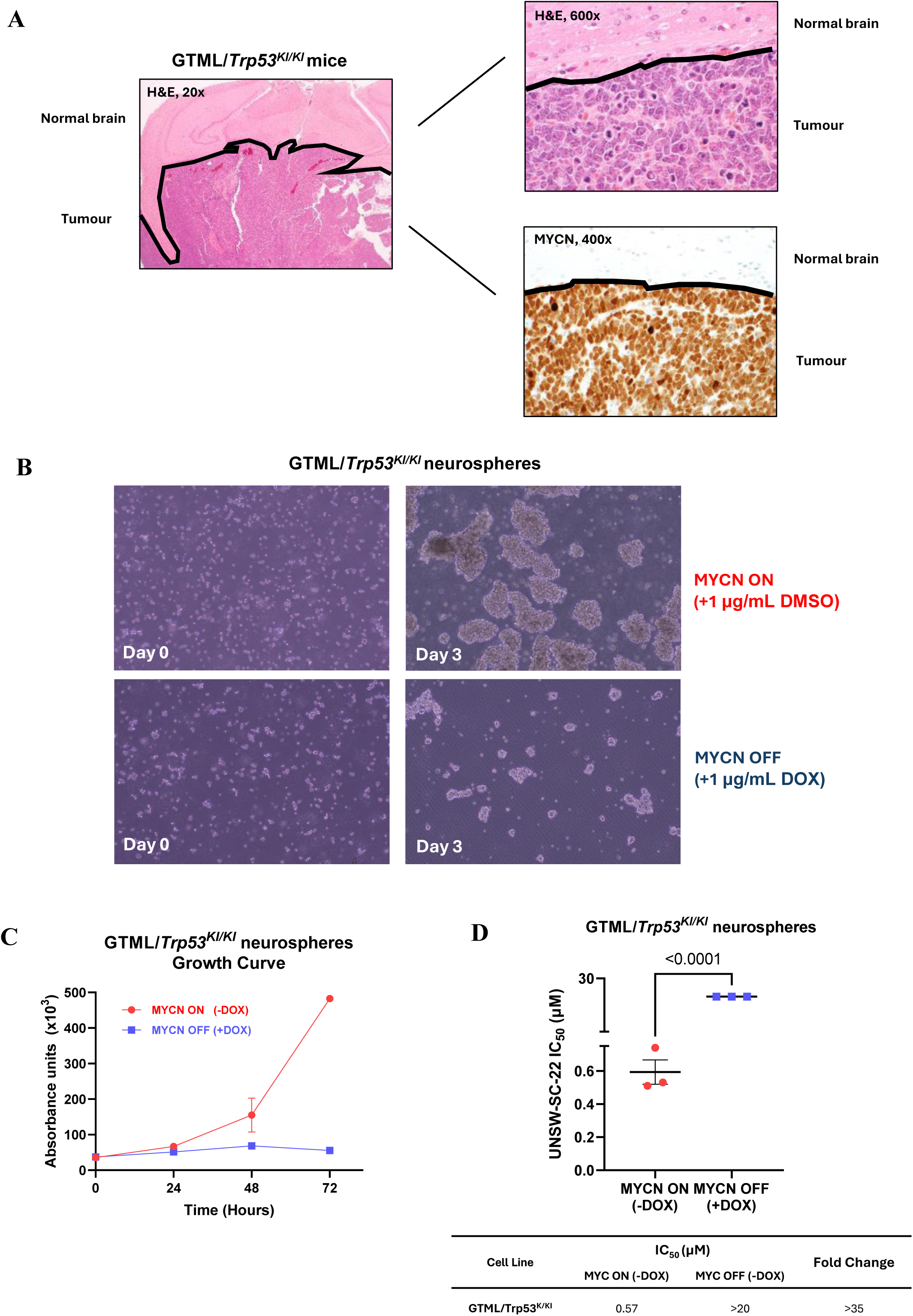
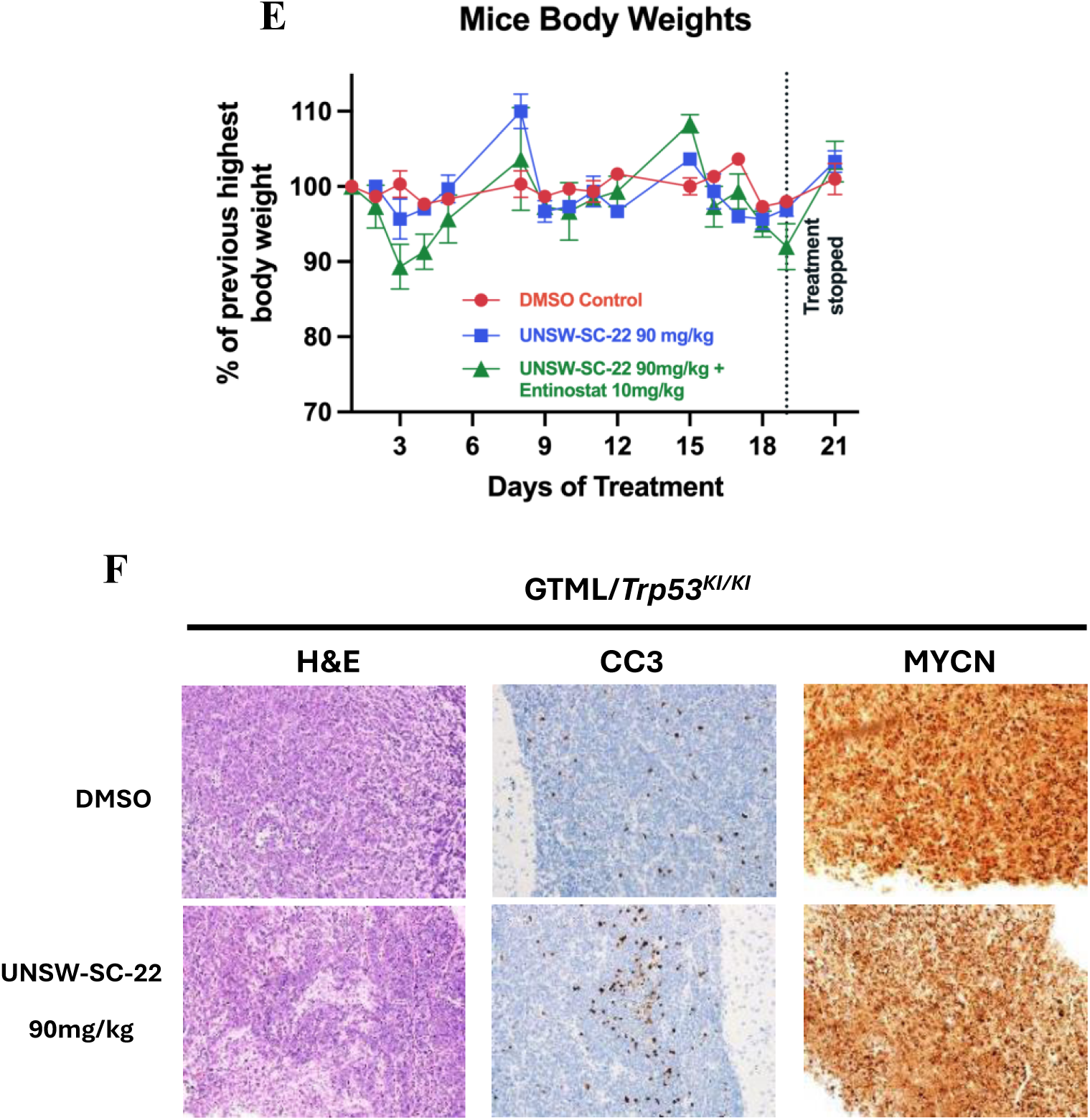
**(A)** Representative images of 50 day old GTML/*Trp53KI/KI* mice brain sections stained with H&E and anti-MYCN at magnification of 20x, 400x or 600x objective lens, and 10x eyepiece lens as indicated. **(B)** Representative images of GTML/*Trp53KI/KI* neurospheres at a magnification of 100x (10x objective lens, 10x eyepiece lens) in the presence of absence of 1μg/μL of doxycycline for 72 hours. **(C)** Growth curve for GTML/*Trp53KI/KI* neurospheres in the presence of absence of 1μg/μL of doxycycline. **(D)** IC50 comparisons for UNSW-SC-22 in GTML/*Trp53KI/KI* neurospheres in the presence or absence of MYCN expression. **(E)** Normalised body weight curves of GTML/*Trp53KI/KI* mice treated with either DMSO control, 90 mg/kg UNSW-SC-22, or a combination with 10 mg/kg entinostat on a treatment schedule of 5 days on, 2 days off for 21 days. Body weights were normalised to the previous highest body weight of each mouse. **(F)** Representative images of treated GTML/*Trp53KI/KI* brain sections at magnification of 200x (20x objective lens, 10x eye piece lens) stained with H&E, anti-Cleaved caspase-3 (CC3), or anti-MYCN. Reported p-values are from one/two-way ANOVA, error bars represent the standard error of the mean (SEM) of at least three independent biological repeats.

## Notes

### Competing Interest Statement

The authors have declared no competing interest.

