## Supplementary methods and materials for "Therapeutic targeting of MYC- and MYCN-driven medulloblastoma with a novel MYC degrader molecule"

### *Study Design*

This study aimed to identify and validate UNSW-SC-22 as a novel MYC degrader for high-risk MB, define its safety and mechanism of action, and evaluate its efficacy alone and in combination with HDAC inhibitors. We first assessed whether UNSW-SC-22 directly engages MYC/MYCN and whether its activity is MYC-dependent, supporting target specificity. Mechanistic studies examined apoptosis, cell-cycle effects, MYC phosphorylation and degradation, and downstream MYC target regulation. We next evaluated BBB penetration and therapeutic efficacy in established MYC- and MYCN-driven MB mouse models. Finally, UNSW-SC-22 was tested in combination with clinically relevant HDAC inhibitors to support translational potential. *In vitro* studies included biological and technical replicates. Animal sample sizes were based on prior experience with similar models, and mice were randomly allocated to treatment groups.

### *Bulk RNA Sequencing*

Total RNA was extracted from D283MED cells treated with vehicle control or UNSW-SC-22 for 24 hours and quantified. For each sample, 1.5 µg RNA was diluted to 20 ng/µL and submitted to the Australian Genome Research Facility (AGRF, Peter MacCallum Cancer Centre) for sequencing. Libraries were prepared using standard Illumina protocols and sequenced on a NovaSeq 6000 at 100 million reads per sample. RNA-seq reads were mapped to the human genome (hg38) using STAR v2.7.5c. Aligned reads were indexed with samtools v1.9, and gene-level counts were generated using HTSeq-count v0.12.4. Downstream analyses were performed in RStudio v2021.09.0. Gene counts were normalised using DESeq2 and filtered using an average TPM threshold of >0.5. Differentially expressed genes were defined as those with an absolute log2 fold change >1 and adjusted p-value <0.05. Data were visualised using ggplot2 and ComplexHeatmap. Pathway enrichment analyses were performed using fgsea v1.20.0 for GSEA with the Molecular Signatures Database, and clusterProfiler v4.2.2.

### Chip-Sequencing Data Analysis

MYC ChIP-seq (GSE143376, h19) bigwig files were obtained from the gene expression omnibus (GEO) for MB cell lines (HDMB03) and patient samples (MB1, MB2 and MB3). The data was then visualised on their respectively aligned genome with the IGV genome browser alongside possible MYC motifs (Ebox) that were generated through CANNTG motif query using the PWMScan tool.

#### *Cycloheximide Chase Assay*

D283MED cells were treated with vehicle control or UNSW-SC-22 for 24 hours, followed by 100 µg/mL cycloheximide (Sigma-Aldrich) for 0, 15, 30, 45 or 60 minutes. Protein lysates were collected at each time point and analysed by western blot to assess MYC stability. Densitometry was normalised to Vinculin and then to the 0-minute time point for each condition. MYC protein half-life was determined using an exponential decay curve in GraphPad Prism 9 from at least three independent biological replicates.

#### *Proteasomal Degradation Assay*

D283MED MB cells were treated with vehicle control or UNSW-SC-22 for 48 hours, and then treated with either DMSO or 30 µM of the proteasome inhibitor MG132 for 6 hours. Protein lysates were then extracted and analysed by western blot. Densitometry was conducted with each sample normalised first to Vinculin as the loading control, and subsequently to their respective vehicle controls.

#### *Flow Cytometric Assays*

Cell cycle distribution was assessed by propidium iodide (PI) staining. D283MED cells were treated with increasing concentrations of UNSW-SC-22 for 24 or 48 hours, harvested, fixed in 80% ethanol, washed, and stained with PI/RNase A in the dark at room temperature. Cells were filtered before analysis on a FACSCanto B. A total of 10,000 cells were acquired per sample, and cell cycle phases were quantified using FlowJo. Apoptosis was assessed using Annexin V/7-AAD staining (BD Biosciences). D283MED cells were treated with increasing concentrations of UNSW-SC-22 for 48 hours, harvested, stained according to the manufacturer's instructions, filtered, and analysed on a FACSCanto B. A total of 10,000 cells were acquired per sample, and apoptotic cells were quantified using FlowJo.

#### *Bioluminescence Imaging*

Tumour growth in orthotopic and transgenic medulloblastoma mouse models was monitored by bioluminescence imaging. Mice were injected intraperitoneally with D-luciferin at 150 mg/kg (GoldBio) and anaesthetised with 2% isoflurane in oxygen. Imaging was performed using an IVIS Spectrum CT with 1.5% maintenance isoflurane. Images were captured from the ventral perspective using a 22.5 cm field of view, small binning, f/8 aperture and auto-exposure of 0.5–5 seconds. Luminescence was measured as photons/sec/cm²/steradian and analysed using Living Image Software (PerkinElmer).

#### *Histology and Immunohistochemistry*

Brain samples were fixed in 10% neutral buffered formalin for 48 hours, then transferred to 70% ethanol at 4°C. Tissues were sagittally sectioned, placed in histology cassettes, processed, embedded, sectioned and stained with haematoxylin and eosin by the Histology and Microscopy Unit (UNSW Sydney). Immunohistochemistry was performed using antibodies against cleaved caspase-3 (Asp175; 1:1000, Cell Signalling Technology), Ki67 (1:250, Merck-Millipore), c-MYC (1:500, Abcam) and MYCN (1:1000, Proteintech). Images were captured using an Olympus BX53 light microscope with a DP-73 camera and cellSens software.
