## Supplementary material for "Therapeutic targeting of MYC- and MYCN-driven medulloblastoma with a novel MYC degrader molecule": Tables

**Table 1. Western Blot Antibodies**

| Target | Vendor/Source | Dilution | Antibody Host |
|---|---|---|---|
| c-MYC | Cell Signalling | 1 : 1000 | Rabbit |
| MELK | Cell Signalling | 1 : 1000 | Rabbit |
| USP1 | Cell Signalling | 1 : 1000 | Rabbit |
| PARP | Cell Signalling | 1 : 1000 | Rabbit |
| Cleaved-caspase-3 | Cell Signalling | 1 : 1000 | Rabbit |
| P21 Waf1/Cip1 | Cell Signalling | 1 : 1000 | Rabbit |
| H2AX | Cell Signalling | 1 : 1000 | Rabbit |
| phospho-H2AX | Cell Signalling | 1 : 500 | Rabbit |
| p53 | Santa Cruz Biotechnology | 1 : 1000 | Mouse |
| MYCN | Santa Cruz Biotechnology | 1 : 1000 | Mouse |
| Vinculin | Sigma-Aldrich | 1 : 5000 | mouse |
| GAPDH | Santa Cruz Biotechnology | 1 : 5000 | mouse |
| β-actin | ThermoFisher | 1 : 10000 | mouse |

**Table 2. Primer sequences used for qRT-PCR:** For each gene, both the forward and reverse sequences are listed.

| Primers | | Sequence |
|---|---|---|
| **MYC** | Forward | ACACTAACATCCCACGCTCT |
| | Reverse | ACAAACCGCATCCTTGTCCT |
| **MYCN** | Forward | CGACCACAAGGCCCTCAGTA |
| | Reverse | CAGCCTTGGTGTTGGAGGAG |
| **MELK** | Forward | CTCAGCCGTGCCCTCC |
| | Reverse | TCCTCCAGGGCAGTACTCAA |
| **USP1** | Forward | AGATGGGCTTGCTGCT |
| | Reverse | ACCGGGACAAAAATATAATACCTGA |
| **TK1** | Forward | GGGGCAGATCCAGGTGATTC |
| | Reverse | AGAATCTGCTCCTTCTCTGTGC |
| **E2F8** | Forward | TGACGAGTCCCGGCTCAG |
| | Reverse | CATCTCCAGATAAGTGTTCGTGT |
| **TOP2A** | Forward | ACCGTCACCATGGAAGTGTC |
| | Reverse | CCATTTCGACCACCTGTCAC |

| | | |
|---|---|---|
| **AURKB** | Forward | GGGAGAGTAGCAGTGCCTTG |
| | Reverse | GAAGTGCCGCCTGGTTAAGA |
| **RRM2** | Forward | GAGGCTACCTATGGTGAACG |
| | Reverse | CCCATCCTCTGATACTCGCC |
| **CHEK1** | Forward | AAACCCCTCAAGAAAGGGGC |
| | Reverse | TGGGGTGCCAAGTAACTGAC |
| **CDC45** | Forward | TGACCACGTGCAATATACGCT |
| | Reverse | GTTCACTCCCAGAGCCACTCC |
| **MCM4** | Forward | ACCCTCAGGACGAAGCCTAT |
| | Reverse | TGGCACTCATCCCCGTAGTA |
| **MCM5** | Forward | GCCTACTGCCGAGTGAAGTGT |
| | Reverse | GGCGTGCTCCGGGTATTTC |
| **B2M** | Forward | ACTGGTCTTTCTATCTCTTGTA |
| | Reverse | TGATGCTGCCTTACATGTCTCG |

**Table 3. Synergism metrics for the combination of UNSW-SC-22 and entinostat in a panel of MB cell lines:** The HSA and BLISS average indicates the averaged HSA and BLISS synergism indices (SI) of the selected dose ratio (UNSW-SC-22: entinostat) identified from the 6x6 matrices. SI > 0 represents synergism, SI = 0 is additive, and SI < 0 indicates antagonism.

| Cell Line | HSA average | BLISS average | UNSW-SC-22 dose (µM) | Entinostat dose (µM) | Ratio |
|---|---|---|---|---|---|
| **D283MED** | 0.25 | 0.14 | 0.5 | 1.25 | 1:2.5 |
| **CHLA-01-MED** | 0.36 | 0.13 | 1 | 0.5 | 1:0.5 |
| **CHLA-01R-MED** | 0.30 | 0.10 | 0.625 | 1.25 | 1:2 |
| **D341MED** | 0.24 | 0.04 | 0.5 | 1.5 | 1:3 |
| **D425MED** | 0.21 | -0.35 | 0.44 | 1.25 | 1:2.8 |

**Table 4. Synergism metrics for the combination of UNSW-SC-22 and quisinostat in a panel of MB cell lines:** The HSA and BLISS average indicates the averaged HSA and BLISS synergism indices (SI) of the selected dose ratio (UNSW-SC-22:quisinostat) identified from

the 6x6 matrices. SI > 0 represents synergism, SI = 0 is additive, and SI < 0 indicates antagonism.

| **Cell Line** | **HSA average** | **BLISS average** | **UNSW-SC-22 dose (µM)** | **Quisinostat dose (µM)** | **Ratio** |
|---|---|---|---|---|---|
| **D283MED** | 0.22 | 0.28 | 0.5 | 0.01 | 1:0.02 |
| **CHLA-01-MED** | 0.21 | 0.02 | 0.1 | 0.01 | 1:0.1 |
| **CHLA-01R-MED** | 0.21 | 0.06 | 0.625 | 0.004 | 1:0.0064 |
| **D341MED** | 0.22 | 0.0095 | 1 | 0.01 | 1:0.01 |
| **D425MED** | 0.28 | 0.05 | 0.44 | 0.003 | 1:6.8 |
